# Behavioral, Anatomical and Genetic Convergence of Affect and Cognition in Superior Frontal Cortex

**DOI:** 10.1101/2020.12.03.401414

**Authors:** Nevena Kraljević, H. Lina Schaare, Simon B. Eickhoff, Peter Kochunov, B.T. Thomas Yeo, Shahrzad Kharabian Masouleh, Sofie L. Valk

## Abstract

Affective experience and cognitive abilities are key human traits that are interrelated in behavior and brain. Individual variation of affective and cognitive traits, as well as brain structure, has been shown to partly underlie genetic effects. However, to what extent affect and cognition have a shared genetic relationship with local brain structure is incompletely understood. Here we studied phenotypic and genetic correlations of cognitive and affective traits in behavior and brain structure (cortical thickness, surface area and subcortical volumes) in the twin-based Human Connectome Project sample (N = 1091). Both affective and cognitive trait scores were highly heritable and showed significant phenotypic correlation on the behavioral level. Cortical thickness in the left superior frontal cortex showed a phenotypic association with both affect and cognition, which was driven by shared genetic effects. Quantitative functional decoding of this region yielded associations with cognitive and emotional functioning. This study provides a multi-level approach to study the association between affect and cognition and suggests a convergence of both in superior frontal cortical thickness.

## 1. Introduction

The human cerebral cortex is implicated in multiple aspects of psychological functions, including cognitive abilities and affective experiences. Psychological traits and neuropsychiatric disorders have been reliably associated with interindividual variation in cortical macrostructure (Thompson et al., 2020). Moreover, variation in macroscale grey matter structure, such as in local cortical thickness and surface area, is strongly driven by hereditary and polygenetic influences (Grasby et al., 2020; Panizzon et al., 2009; Winkler et al., 2010). Affective and cognitive traits have also been shown to underlie genetic effects (Davies et al., 2011; Okbay et al., 2016; Zheng et al., 2016). However, to which degree trait affect, cognition, and brain structure share a genetic basis is incompletely understood.

Twin-based genetic studies have previously been conducted to assess heredity of habitual (i.e. trait) cognitive and affective processes. Such pedigree-based designs allow the assessment of genetic effects on a given phenotype by comparing monozygotic twins with other sibships and unrelated individuals (Almasy et al., 1997; Almasy and Blangero, 1998). Twin-based studies showed that cognitive abilities are largely influenced by genetic effects (Bartels et al., 2002; Davies et al., 2011; Kan et al., 2013; van Soelen et al., 2011; Wainwright et al., 2005). Affective traits have been investigated in genetic studies by using measures of positive and negative affect, as well as subjective well-being (Diener and Emmons, 1984; Lykken and Tellegen, 1996; Russell and Carroll, 1999; Salsman et al., 2013; Watson and Tellegen, 1985). These studies repeatedly found trait negative affect to be heritable, while trait positive affect was not related to genetic effects (Baker et al., 1992; Zheng et al., 2016). Conversely, Lykken and Tellegen (1996) showed that individual differences in subjective well-being were partly explained by genetic variation in several thousand middle-aged twins. Along this line, Genome-Wide Association Studies (GWAS) reported various loci associated with subjective well-being (Okbay et al., 2016). These results emphasize a genetic basis of the variation in both cognition and some affective experiences.

Human brain structure also underlies genetic influences, as brain volume, cortical surface area and thickness have been found to be strongly heritable and to show a polygenetic architecture (Brouwer et al., 2014; Grasby et al., 2020; Panizzon et al., 2009; Winkler et al., 2010). Neuroimaging genetics studies demonstrated that associations between cognition and cortical thickness can be explained by shared genetic effects (Brans et al., 2010; Desrivières et al., 2015; Joshi et al., 2011; Shaw et al., 2006). Gene enrichment studies have further associated subjective well-being with differential gene expression in the hippocampal subiculum and with GABAergic interneurons, suggesting a genetic link between brain structure and affective traits (Baselmans et al., 2019). Moreover, shared genetic effects have been found to drive the association between neuroticism – a personality trait closely linked to negative affect – and surface area in the right medial frontal cortex (Valk et al., 2020).

Classically considered to be distinct entities, most neuroimaging studies to date investigated neural correlates of cognition and affect as separate constructs yielding inconclusive results. There is a multitude of evidence which suggests that general cognitive abilities are positively correlated with greater brain volume across the human lifespan (Oschwald et al., 2019). Using surface-based measures, individual differences in cognition have been related to prefrontal and parietal cortical thickness, though often with contradictory outcomes: Both positive correlations between cortical thickness and cognitive abilities (Karama et al., 2009; Narr et al., 2006; Shaw et al., 2006, Bajaj et al., 2018, Hanford et al., 2019), as well as negative correlations (Goh et al., 2011; Salat et al., 2002; Sowell et al., 2001; Van Petten et al., 2004) have been reported. With respect to affective traits, neuroimaging studies have repeatedly shown that state and trait emotional processes correlate with activity, connectivity and anatomy of the brain (Atkinson et al., 2007; Brierley et al., 2004; LaBar et al., 1995; Lindquist et al., 2012; Rohr et al., 2015; Tsuchiya et al., 2009).

Recent theories emphasize the interplay and shared mechanisms of emotions and cognition in the human brain (Barrett, 2016; Pessoa, 2008): Emotions can both facilitate and impede cognitive function, depending on the context (Dolcos and Denkova, 2014; Okon-Singer et al., 2015). At the same time, cognitive processes are inherent in most aspects of emotional experience and regulation (Ochsner and Gross, 2005; Pessoa, 2008). This behavioral association is mirrored by overlapping brain networks associated with emotions (Barrett, 2016; Khalsa et al., 2018) and cognitive control (Langner et al., 2018; Pessoa, 2008). Thus, affective experience and cognitive abilities are inherently coupled in the human brain (Barrett, 2016; Pessoa, 2008). Yet, it remains unclear if this coupling reflects in neuroanatomical correlates and if cognition and affect share a genetic basis.

In sum, 1) cognitive abilities and their relation with brain structure are under genetic control; 2) affective experience is associated with brain function and structure and may also be driven by genetic factors, depending on the affect measurement; 3) there is a complex interrelation between affect and cognition in behavior and brain. However, whether cognition and trait affect have a shared genetic relation to brain structure is not known to date. We studied the relationship of cognition and affect in behavior and local brain structure and evaluated whether cognitive abilities and trait affective self-reports have a shared genetic basis in both behavior and brain. First, we evaluated the relation of cognition and trait affect on the behavioral level by conducting phenotypic correlation, as well as heritability analyses and genetic correlation in a large sample of healthy twins. Next, we assessed the relationship between cognition and affective traits in relation to cortical thickness, subcortical volumes and cortical surface area, to evaluate whether cognition and affect share phenotypic and genetic associations with local brain structure. We expected to observe that correlations of cognition and affect with the brain can be explained by shared genetic effects.

## 2. Materials and methods

### 2.1. Participants

The Human Connectome Project (HCP) is a publicly available data base. In this study, the Young Adult Pool was used, which comprised 1206 healthy individuals (656 women, mean age = 28.8 years, standard deviation (SD) = 3.7, range = 22-37 years). In total, there were 292 monozygotic (MZ) twins, 323 dizygotic (DZ) twins, and 586 singletons (additionally 5 missing values in zygosity information). After exclusion of individuals without brain structural (N = 93) or behavioral data (N = 20) relevant to this study, and participants with corrupted brain data (N = 4), our final sample comprised 1091 individuals, of which 592 were women. This sample included 274 MZ twins, 288 DZ twins, and 525 singletons (additionally 4 missing values in zygosity information). Its mean age, standard deviation and range remained the same as for the total HCP sample.

### 2.2. Ethics statement

Analysis of the HCP data has been approved through the local ethics committee of the University of Düsseldorf, Germany.

### 2.3. Data/code availability statement

To ensure reproducibility of this study using unrestricted and restricted data of the publicly available HCP dataset (www.humanconnectome.org), the code that has been used in our analyses can be found here: https://github.com/CNG-LAB/affect_cognition. As specified in the [HCP Restricted Data Use Terms](https://www.humanconnectome.org/study/hcp-young-adult/document/wu-minn-hcp-consortium-restricted-data-use-terms), investigator-assigned IDs of included participants will be shared upon publication of the study.

### 2.4. Behavioral measures

The cognitive and affective measures used in this study were selected in the data base of the HCP and derived from the National Institute of Health (NIH) toolbox for Assessment of Neurological and Behavioral Function^®^ (neuroscienceblueprint.nih.gov). Composite scores from the cognition and emotion categories were used, while one category comprised several sub-domains (Table 1).

**Table 1.** Behavioral scores. Overview of composition of behavioral variables.

| Category | Domain | Sub-domain | Test |
| --- | --- | --- | --- |
| Cognition | Fluid cognition | Executive function – cognitive flexibility | Dimensional Change Card Sorting (DCCS) |
|  |  | Executive function – Inhibition and attention | Flanker |
|  |  | Episodic memory | Picture Sequence Memory |
|  |  | Processing speed | Pattern Comparison |
|  |  | Working memory | List Sorting |
|  | Crystallized cognition | Language | Picture Vocabulary |
|  |  |  | Reading Recognition |
| Affect | Positive affect/ psychological well-being | Life satisfaction | Self-report |
|  |  | Meaning and purpose |  |
|  |  | Positive affect |  |
|  | Negative affect | Anger-affect | Self-report |
|  |  | Anger-hostility |  |
|  |  | Fear-affect |  |
|  |  | Perceived stress |  |
|  |  | Sadness |  |

#### 2.4.1. Cognition

The cognitive function composite score (total cognition) was assessed by averaging the fluid cognition composite score (fluid cognition) and the crystallized cognition composite score (crystallized cognition). As illustrated in Table 1, the fluid cognition score was obtained by averaging the scores of the Dimensional Change Card Sort Test, Flanker, Picture Sequence Memory, List Sorting, and Pattern Comparison measures. That is, fluid cognition is the combination of scores of executive function, inhibition and attention, episodic memory, working memory, and processing speed (Akshoomoff et al., 2013). The crystallized cognition score was obtained by averaging the scores of Picture Vocabulary and Oral Reading Recognition measures. That is, crystallized cognition consists of language in the sense of translation of thought into symbols and deriving meaning from text, as a reflection of past learning experiences (Akshoomoff et al., 2013; Gershon et al., 2013).

As cognition can be both conceived as a general factor (G), but at the same time crystallized and fluid cognition are differentiable, we investigated the general cognitive score (total cognition), as well as fluid and crystallized cognition.

#### 2.4.2. Affect

To examine trait affect in this study, a composite measure of general affect was used, consisting of both positive and negative affect. Trait affect can be sub-divided into positive and negative traits, which are separable constructs that may be represented as one bipolar scale or two unipolar scales (Diener and Emmons, 1984; Russell and Carroll, 1999; Salsman et al., 2013; Watson and Tellegen, 1985). Positive affect, also known as psychological well-being, is characterized by the experience of pleasant feelings, such as happiness, serenity and cognitive engagement (Diener and Emmons, 1984; Salsman et al., 2014, 2013). We composed the construct of positive affect by averaging the scores from the sub-domains of life satisfaction, meaning and purpose, and positive affect (Table 1, Table 2). Negative affect comprises three principal negative emotions: anger, fear, and sadness (Pilkonis et al., 2013; Salsman et al., 2013). It was composed by the average of anger (anger-affect, hostility), sadness, fear-affect, and perceived stress (Table 1, Table 2). All affective domains were obtained using the NIH toolbox with a written self-report (Pilkonis et al., 2013; Salsman et al., 2014, 2013).

**Table 2.** Behavioral variables. Mean, standard deviation, as well as minimum and maximum of each variable included in our analyses.

| Variables | Mean | SD | Min | Max |
| --- | --- | --- | --- | --- |
| Total Cognition | 121.8 | 14.6 | 84.6 | 153.4 |
| Fluid Cognition | 115.0 | 11.6 | 84.5 | 145.2 |
| Crystallized Cognition | 117.7 | 9.9 | 90.4 | 154.0 |
| Positive Affect | 52.1 | 7.2 | 27.1 | 72.6 |
| Negative Affect | 48.7 | 6.8 | 30.9 | 78.8 |
| Mean Affect | 1.7 | 6.2 | -19.6 | 20.8 |

As the positive and negative affect scores used in this study showed high intercorrelations (R = −0.6, see Figure 1C and Supplementary Figure 1), we created a composite score of mean affect by reversing the negative affect score and averaging it with the positive affect score. This enabled us to investigate positive and negative affect as separate entities, on the one hand, as well as a general estimate of mean affect that integrates both negative and positive emotions, on the other hand.

**Figure 1.**
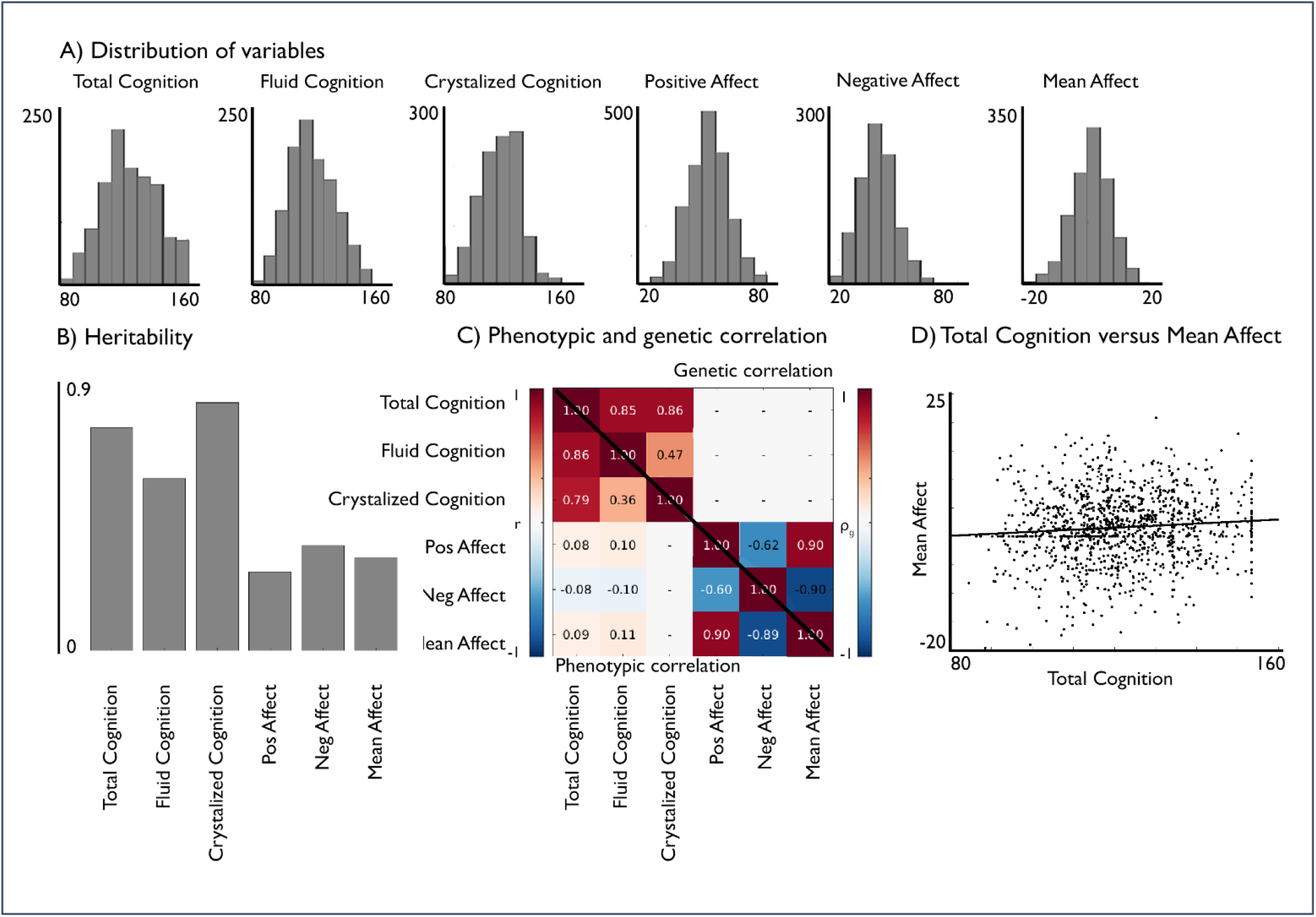
Phenotypic and genetic relation of cognition and affect. A) Distribution of variables; B) Heritability; C) bottom triangle: behavioral correlation (FDRq<0.05) and upper triangle: genetic correlation of the cognitive and affective scores (FDRq <0.05).

### 2.5. Structural imaging processing

MRI protocols of HCP were previously described in detail (Glasser et al., 2013; Van Essen et al., 2013). In short, MRI data used in the study was acquired on the HCP’s custom 3T Siemens Skyra scanner equipped with a 32-channel head coil. Two T1-weighted (T1w) images with identical parameters were acquired using a 3D-MPRAGE sequence (0.7 mm isotropic voxels, matrix = 320 × 320, 256 sagittal slices, TR = 2,400 ms, TE = 2.14 ms, TI = 1,000 ms, flip angle = 8°, iPAT = 2). Two T2w images were acquired using a 3D T2-SPACE sequence with identical geometry (TR = 3,200 ms, TE = 565 ms, variable flip angle, iPAT = 2). T1w and T2w scans were acquired on the same day. The pipeline used to obtain the FreeSurfer segmentation is described in detail in a previous article (Glasser et al., 2013) and is recommended for the HCP-data. The pre-processing steps included co-registration of T1w and T2w scans, B1 (bias field) correction, and segmentation and surface reconstruction using FreeSurfer version 5.3-HCP to estimate brain volumes, cortical thickness and surface area. We also derived eight bilateral subcortical volumes from FreeSurfer’s automatic subcortical segmentation (Fischl et al., 2002) to evaluate their phenotypic and genetic correlation with behavioral traits.

### 2.6. Cortical morphological measures

For analyses including cortical structure, we used a parcellation scheme (Schaefer et al., 2018) based on the combination of local gradient and global similarity approaches using a gradient-weighted Markov Random Field model. Using compressed features of structural MRI has been suggested to both improve signal-to-noise of brain measures (cf. Eickhoff et al., 2018; Genon et al., 2018) and optimize analysis scalability. The Schaefer parcellation has been extensively evaluated with regards to stability and convergence with histological mapping and alternative parcellations (Schaefer et al., 2018). In the context of the current study, we focused on the granularity of 200 parcels from the 7-network solution. In order to improve signal-to-noise ratio and analysis speed, we opted to average unsmoothed structural data within each parcel. Thus, cortical thickness of each parcel was estimated as the trimmed mean (10 percent trim) and parcel-wise surface area was computed as the sum of vertex-wise area per parcel.

### 2.7. Phenotypic correlation analyses

Phenotypic correlations between cognitive and affective traits were assessed by cross-correlating the normalized behavioral measures, controlling for effects of age, sex and their interaction, using multiple linear regression models.

Phenotypic analyses between behavioral traits and local brain structure were carried out per parcel of cortical thickness and surface area, as well as per volume of subcortical structures.

Each brain modality was predicted by cognition and affect, respectively, using multiple linear regression models while controlling for age, sex, age × sex interaction, age^2^, age^2^ × sex interaction, as well as global thickness (mean cortical thickness) effects when investigating cortical thickness and intracranial volume (ICV) when assessing surface area and subcortical volumes. As in previous work (Bernhardt et al., 2014; Valk et al., 2016a, 2016b), we used SurfStat for Matlab [R2020a, The Mathworks, Natick, MA] (Worsley et al., 2009) to conduct the statistical comparisons.

Results of all phenotypic correlations were corrected for multiple comparisons using Benjamini-Hochberg false discovery rate (FDRq<0.05) (Benjamini and Hochberg, 1995). We displayed significant brain associations on the cortical surface.

### 2.8. Heritability and genetic correlation analyses

To investigate the heritability and genetic correlation of brain structure and behavioral traits, we analyzed 200 parcels of cortical thickness, surface area, and subcortical volumes, as well as cognitive and affective trait scores of each participant in twin-based heritability and genetic correlation analyses. As in previous studies (Glahn et al., 2010), the quantitative genetic analyses were conducted using Sequential Oligogenic Linkage Analysis Routines (SOLAR) (Almasy and Blangero, 1998; Kochunov et al., 2019). SOLAR uses maximum likelihood variance-decomposition methods to determine the relative importance of familial and environmental influences on a phenotype by modeling the covariance among family members as a function of genetic proximity. This approach can handle pedigrees of arbitrary size and complexity and thus, is optimally efficient with regard to extracting maximal genetic information (Almasy and Blangero, 1998). To ensure that our traits, behavioral as well as of brain structure, were conform to the assumptions of normality, an inverse normal transformation was applied for all behavioral and neuroimaging traits (Glahn et al., 2010). Heritability (*h*^2^) represents the proportion of the phenotypic variance 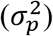 accounted for by the total additive genetic variance 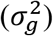, i.e., 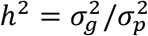. Phenotypes exhibiting stronger covariances between genetically more similar individuals than between genetically less similar individuals have higher heritability. Heritability analyses were conducted with simultaneous estimation for the effects of potential covariates. We thus included the same covariates as in our phenotypic correlation analysis including age, sex, age × sex interaction, age^2^, age^2^ × sex interaction, as well as global thickness when investigating cortical thickness and ICV when assessing surface area and subcortical volumes. Heritability estimates were corrected for multiple comparisons by Benjamini-Hochberg FDRq<0.05 (Benjamini and Hochberg, 1995).

To determine if variations in cognition or affect and brain structure were influenced by the same genetic factors, genetic correlation analyses were conducted. More formally, bivariate polygenic analyses were performed to estimate genetic (*ρ_g_*) and environmental (*ρ_e_*) correlations, based on the phenotypic correlation (*ρ_p_*), between brain structure and behavioral traits with the following formula: 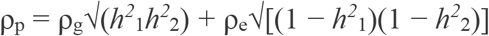, where *h*^2^_1_ and *h*^2^_2_ are the heritability estimates of the parcel-based brain structural measures and the respective behavioral traits. The significance of these correlations was tested by comparing the log likelihood for two restricted models (with either ρg or ρe constrained to be equal to 0) against the log likelihood for the model in which these parameters were estimated. A significant genetic correlation (corrected for multiple comparisons using Benjamini-Hochberg FDR (FDRq<0.05) (Benjamini and Hochberg, 1995)) indicates that (a proportion of) both phenotypes are influenced by the same gene or set of genes (Almasy et al., 1997). In all genetic correlation analyses, we controlled for the same covariates as in phenotypic and heritability analyses, including age, sex, age × sex interaction, age^2^, age^2^ × sex interaction and global cortical thickness (for cortical thickness analyses) or total intracranial volume (for surface area and subcortical volume analyses).

### 2.9. Functional decoding

Parcels that were significantly correlated with both cognition and affect and shared genetic variance, were functionally characterized using the Behavioral Domain meta-data from the BrainMap meta-analysis database (http://www.brainmap.org, Laird et al., 2011, 2009). The BrainMap database enables the decoding of functions associated with specific brain regions. To investigate potential functional processes associated with parcels linked to both affect and cognition, we used the volumetric counterparts of the surface-based parcels as defined by Schaefer et al and available online (Schaefer et al., 2018, https://github.com/ThomasYeoLab/CBIG/tree/master/stable_projects/brain_parcellation/Schaefer2018_LocalGlobal/Parcellations). Thus, although our empirical analysis focused on surface-based cortical data, functional decoding could be performed in volume space. In particular, we identified those meta-data labels (describing the computed contrast [behavioral domain as well as paradigm]) that were significantly more likely than chance to result in activation of a given parcel (Fox et al., 2014; Genon et al., 2018; Nostro et al., 2017). That is, functions were attributed to the parcels by quantitatively determining which types of experiments were associated with activation in the respective parcel region. Significance was established using a binomial test (q<0.05, corrected for multiple comparisons using FDR) and we report results of both forward and reverse inference analyses.

## 3. Results

### 3.1. Heritability, phenotypic and genetic correlation of cognition and affect (Figure 1)

First, we evaluated the phenotypic associations between cognitive test scores and affective self-report scores (**Figure 1, Supplementary Figure 1**). Both composite scores (total cognition and mean affect), as well as their sub-domains (fluid cognition, crystallized cognition, positive affect, and negative affect), were normally distributed (**Figure 1A**). We observed high phenotypic interrelationships between the respective sub-tests of both cognitive and affective domains (**Supplementary Figure 1**), supporting the use of the composite scores total cognition and mean affect as proxies for the constructs of cognition and affect, respectively. As expected, cognitive scores were all positively associated among each other, while positive affect correlated positively, and negative affect correlated negatively to mean affect. We observed that mean affect had a positive phenotypic relationship with total cognition (β = 0.09, FDRq < 0.05). Similarly, positive affect was positively associated with total (β = 0.08) and fluid (β = 0.10) cognitive abilities, whereas negative affect was negatively associated with total (β = −0.08) and fluid (β = −0.10) cognitive abilities (all FDRq < 0.05; **Figure 1C**, lower triangle).

The heritability analysis revealed that all observed construct scores were heritable: total cognition (h^2^= 0.75, p < 0.001), which is a combination of fluid (h^2^= 0.58, p < 0.001) and crystallized (h^2^= 0.83, p < 0.001) cognition, as well as mean affect (h^2^= 0.31, p < 0.001), which is the signed average of positive (h^2^= 0.27, p < 0.001) and negative (h^2^= 0.36, p < 0.001) affect (**Figure 1B**).

Next, we evaluated the genetic correlation between cognitive and affective scores. A strong positive genetic correlation between fluid and crystallized cognition (ρ_g_ = 0.47, FDRq<0.05) and a negative genetic correlation between positive and negative affect (ρ_g_ = −0.62, FDRq<0.05) was found (**Figure 1C**, upper triangle), suggesting that both sub-domain-sets reflect partly overlapping genetic mechanisms. We did not observe a significant genetic correlation between total cognition and mean affect (FDRq > 0.05). At trend level, we observed genetic correlations between fluid cognition and mean affect (ρ_g_ = 0.23, p<0.03), and between fluid cognition and positive affect (ρ_g_ = 0.28, p<0.02). In the following, we focus on reporting the results for the composite scores of total cognition and mean affect which sufficiently capture phenotypic and genetic variance of the constructs of cognition and affect. To additionally allow for more detailed assessment, we report associations with the sub-domain measures in the supplementary materials.

### 3.2. Phenotypic association between cognition and local brain anatomy (Figure 2)

To evaluate the phenotypic association of cognition and affect with brain anatomy, we first evaluated the correlation between cognition and local cortical thickness, while controlling for global thickness. We observed positive associations between thickness and total cognition in bilateral insula, bilateral cuneus, bilateral sensorimotor regions, left middle temporal gyrus, and right middle cingulate; whereas bilateral frontal regions and left parietal showed a negative relation with total cognitive score (**Figure 2A**). Conversely, surface area showed only positive, but not negative, associations with cognitive scores. Total cognition was associated with local surface area in bilateral occipital areas, temporal poles, sensorimotor cortices, and lateral and orbital frontal cortices, as well as left anterior cingulate and right posterior-mid cingulate cortex (**Figure 2B**). With regards to subcortical volumes, we observed a significant positive effect between total cognition and left hippocampal volume (**Figure 2C**). Fluid and crystallized cognition sub-scores showed similar associations with local brain structure and volume (**Supplementary Tables 1-3, 7-12**).

**Figure 2.**
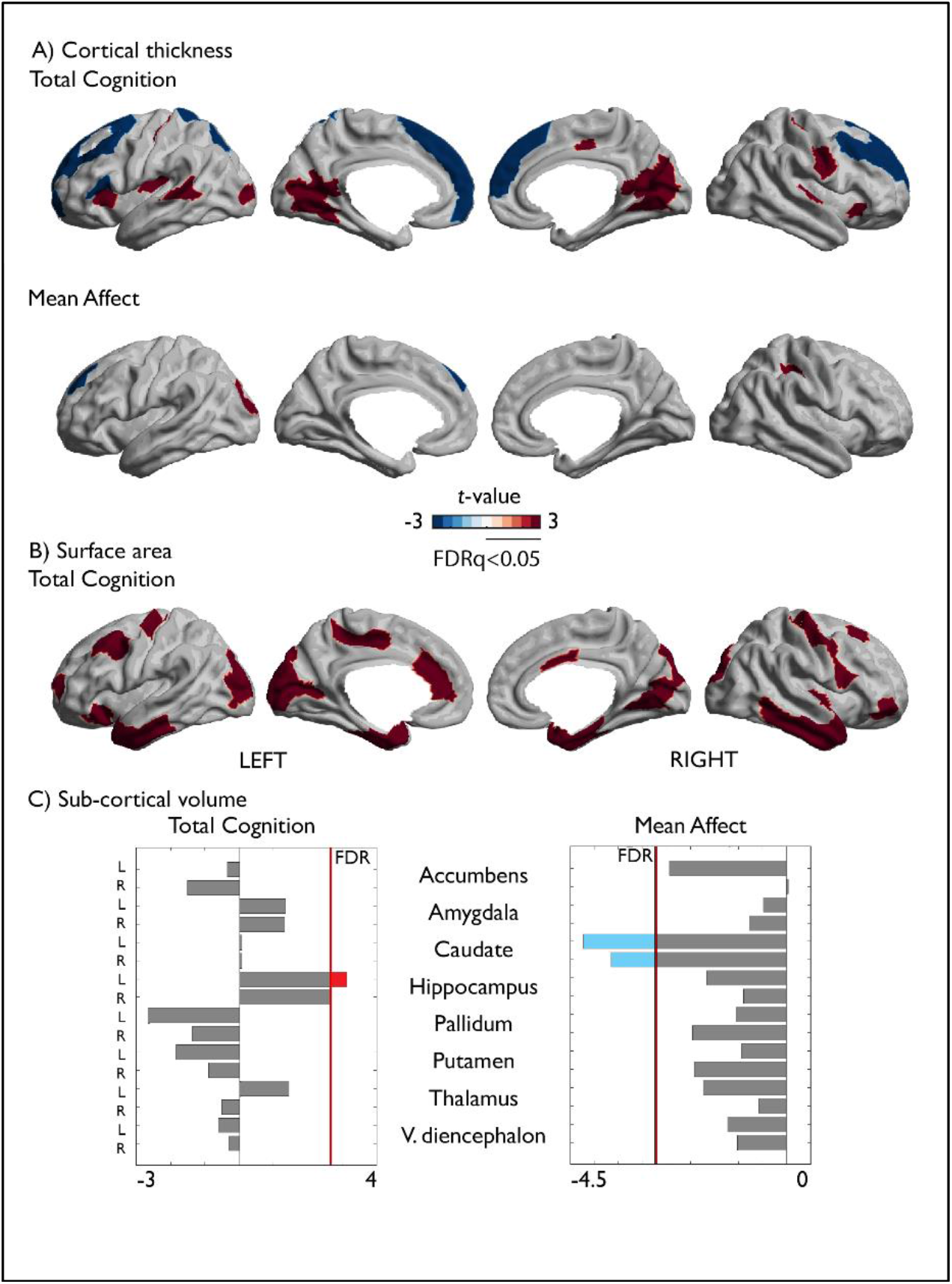
Associations between cognition, affect and local brain structure. A) Correlation between total cognition and local cortical thickness; Second row: Correlation between mean affect and local cortical thickness. B) Correlation between total cognition and local surface area. Associations between surface area and affect were not significant. C) Correlations between cognition / affect and sub-cortical regions volumes. Red indicates a positive association, and blue a negative association between cognition / affect and local brain structure. Only FDRq<0.05 corrected findings are depicted.

### 3.3. Phenotypic association between affect and local brain anatomy (Figure 2)

Next, we evaluated the association between affect and local brain structure. We found that affect measures showed significant associations with local cortical thickness and subcortical volumes, but not with local surface area. Mean affect was associated with cortical thickness negatively in left superior frontal cortex and positively in left occipital cortex, as well as right parietal cortex (**Figure 2A, Supplementary Table 4**). In addition, bilateral caudate volume was significantly associated with mean affect (**Figure 2C, Supplementary Table 13**).

### 3.4. Genetic correlation of cognition and affect with local brain structure (Figure 3)

To assess if the correlation between cognitive and affective traits on the one hand and local brain structure on the other is driven by shared genetic effects, we performed genetic correlation analyses through a bivariate polygenetic analysis. Both local thickness and surface area were heritable in our sample (cortical thickness h^2^=mean±sd: 0.35±0.11 and surface area: h^2^=0.42±0.13), as were subcortical volumes (h^2^=0.68±0.10, **Supplementary Figure 4**, **Supplementary Tables 21-23**).

There was a strong overlap between phenotypic correlations and genetic correlations. 34 out of 37 phenotypic correlations between total cognition and local cortical thickness could be attributed to genetic effects (FDRq<0.05, **Figure 3A, Supplementary Table 16**) and genetic correlation patterns largely mirrored phenotypic associations between cognitive scores and local cortical thickness (see also **Supplementary Figure 5** and **Supplementary Table 18**). Similarly, the phenotypic associations between cognitive scores and subcortical volumes were genetically driven (**Supplementary Table 20**). Furthermore, the associations between local surface area and cognitive scores were largely driven by shared genetic factors. Here, 29 out of 42 phenotypic associations between total cognition and local surface area were driven by shared genetic factors (**Figure 3B, Supplementary Table 17; Supplementary Figure 6** and **Supplementary Table 19** for sub-score results). Among the associations with mean affect, thickness of left superior frontal cortex (ρ_g_ = −0.480, p=0.000, **Figure 3A, Supplementary Table 16**) and bilateral caudate volumes (left: ρ_g_ = −0.28, p=0.001, right: ρ_g_ = −0.28, p=0.001, **Supplementary Table 20**) were also driven by shared genetic factors.

**Figure 3.**
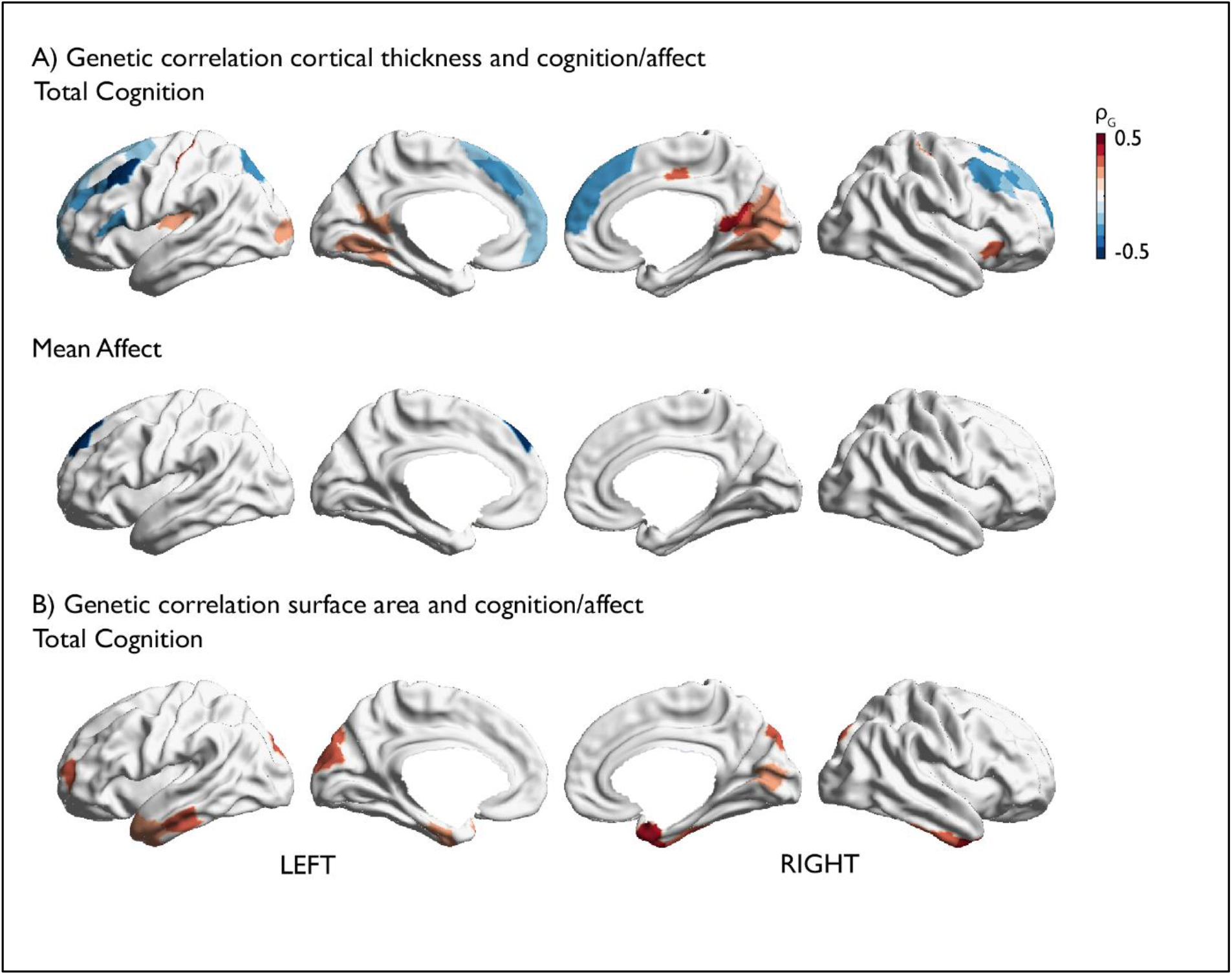
Whole-brain genetic correlation between local cortical structure and cognition or affect. A) Results for cortical thickness. B) Results for surface area. Positive correlation is depicted in red, negative in blue. Only FDRq<0.05 corrected findings are depicted.

### 3.5. Shared brain basis between cognitive and affective tendencies (Figure 4)

Last, we evaluated whether cognitive and affective traits also showed an overlapping relationship to local brain structure. Both cognition and affect scores had an association (FDRq < 0.05) with thickness in the left superior frontal cortex. In both measures, these effects were driven by shared genetic factors (**Supplementary Tables 16-17**). We performed functional decoding to further quantify the functional processes associated with this region and found this region to be involved in cognitive and socio-cognitive processes, as well as emotional processes (valence and negative emotions) and action inhibition (**Figure 4**).

**Figure 4.**
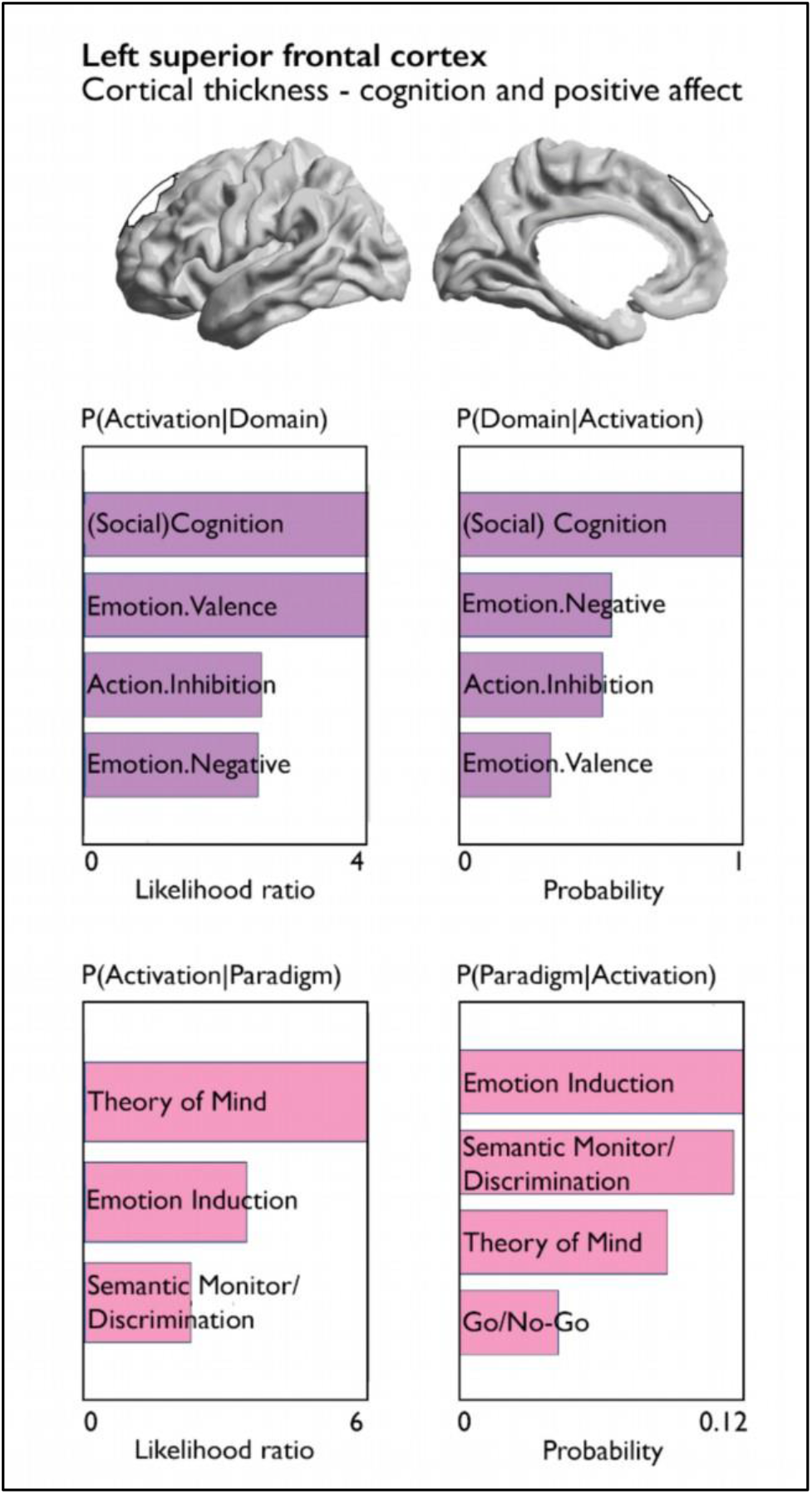
Quantitative functional decoding of region showing association with both cognition and affect. Both forward inference and reverse inference of activation-domain and paradigm-domain contrasts are reported for the left superior frontal cortex which showed evidence of shared phenotypic and genetic association for cognition and affect.

## 4. Discussion

We evaluated shared behavioral, genetic and brain structural factors of cognitive and affective traits. We found that both cognitive and affective traits were heritable and observed significant genetic correlation between fluid cognition (but not total cognition) and trait affect. Following, we assessed the correlation between cognitive and affective traits on the one hand, and macroscale brain anatomy on the other. Whereas cognition had widespread associations with local cortical thickness and surface area, trait affect showed only sparse associations. We found that most phenotypic behavior-brain associations were driven by shared genetic effects. Finally, we evaluated whether total cognition and mean affect were embedded in a common brain structural correlate and found that both measures showed a shared phenotypic and genetic association with cortical thickness of left superior frontal cortex. Quantitative functional decoding further indicated that this region is involved in both cognitive and emotional functioning.

### 4.1. Heritability and genetic correlations of cognition, affect and brain structure

Complementing previous studies on affect, cognition, and macroscale brain anatomy, we interrogated the genetic basis of cognition and affect using twin-based genetic approaches. We observed a moderate to strong heritability of cognitive scores (h^2^ = 0.6-0.8), which is in line with previous work: In childhood and adolescence – depending on measurement and cohort −70 to 80 % of the variance in cognitive ability is estimated to be accounted for by genetic factors (Bartels et al., 2002; van Soelen et al., 2011; Wainwright et al., 2005). Using GWAS of adult samples, Davies et al. (2011) observed that 40% of the variation in crystallized-type intelligence and 51% of the variation in fluid-type intelligence between individuals is accounted for by genetic variants. Notably, crystallized cognition was observed to be more heritable than fluid cognition in the current sample. These findings are in line with a large meta-analysis assessing the heritability of cognitive traits based on their cultural load, where traits with higher cultural load were shown to be more heritable (Kan et al., 2013). Traits that we summarized as crystallized cognition were attributed a higher cultural load in Kan et al.’s study. This indicates that the known cultural and educational homogeneity of the HCP sample may have led to a high estimated heritability of crystallized cognition.

However, our findings on the heritability of affective self-reports were less strong. Previous work in a twin sample by Baker et al. revealed a strong heritability for negative affect, but none for positive affect (Baker et al., 1992), which was conceptually replicated by Zheng et al. (2016). In addition, angry temperament has been associated with genetic processes involved in memory and learning (Mick et al., 2014). Another twin study by Lykken and Tellegen (1996) found the heritability for subjective well-being to be 44-52%, which appears to be higher than the heritability scores we observed for affect measures. However, few studies to date have assessed the heritability of both cognitive and affective traits in the same cohort. Our observations suggest that inter-individual variance in cognition is more robustly explained by genetic factors, than in affect. This could be related to the challenge to quantify individual difference of affective traits in self-reports, which show weaker convergent validity, as opposed to tests for cognitive assessments (Heaton et al., 2014; Salsman et al., 2013).

Moreover, we replicated previous results that showed heritability of surface area and cortical thickness, which further indicated that both variance in cortical thickness and surface area are partly driven by largely non-overlapping genetic factors (Brouwer et al., 2014; Grasby et al., 2020; Panizzon et al., 2009; Winkler et al., 2010).

Extending previous work, we also observed strong genetic correlations between total cognition and local cortical structure which indicates that the majority of phenotypic associations between total cognition and cortical thickness and surface area, respectively, could be associated with shared genetic factors (Brouwer et al., 2014; Grasby et al., 2020; Toga and Thompson, 2005). Affect was phenotypically correlated with superior frontal thickness and genetic correlation analyses yielded that this association was driven by shared genetic effects. These results are in line with recent work implicating various genetic loci with well-being which showed significant enrichment for GABAergic interneurons sampled from hippocampus and prefrontal cortex (Baselmans et al., 2019; Okbay et al., 2016). However, we did not observe genetic associations between affect and hippocampal volumes in this sample. Moreover, affect was genetically correlated with bilateral caudate volumes, which have been shown to share a genetic basis with neuropsychiatric health traits (Satizabal et al., 2019; Zhao et al., 2019).

### 4.2. Shared basis of cognition and affect in behavior, genetics and superior frontal cortex thickness

We combined behavioral and brain imaging approaches to study the association between cognition and trait affect. Scores for mean total cognition and affect showed a positive association at the behavioral level, highlighting the synergy of cognitive and affective traits. Previous work has suggested positive affect might have a motivating role in enhancing cognitive flexibility (Ashby et al., 1999; Fredrickson, 2001; Liu and Wang, 2014). In turn, cognitive control is a core feature of successful emotion regulation (Engen and Anderson, 2018; Ochsner and Gross, 2005) and contributes to psychological well-being over the lifespan (Mather and Carstensen, 2005).

Recent work indicated a shared genetic basis between local brain structure and complex behavioral traits (Grasby et al., 2020; Zhao et al., 2019). In line with this work, our study demonstrated that cognition and trait affect have a shared phenotypic and genetic relationship with cortical thickness in left superior frontal cortex. The superior frontal gyrus – which includes the dorsolateral prefrontal cortex – has been classically considered a core region for higher cognitive functions, including attention, working memory and cognitive control (Boisgueheneuc et al., 2006; Corbetta and Shulman, 2002). Yet, a growing body of research has highlighted its involvement in socio-emotional processes, such as motivated behavior and emotion regulation (Engen and Anderson, 2018; Frank et al., 2014; Okon-Singer et al., 2015).

Left superior frontal gyrus has also been implicated in self-awareness and introspection (Goldberg et al., 2006), as well as in psychiatric disorders of self-awareness, such as schizophrenia (Lee et al., 2016). Indeed, left superior frontal thickness has been shown to be modulated by schizophrenia-associated genetic variants, suggesting a shared genetic basis of schizophrenia-associated brain regions and the neurocognitive symptoms characterizing the disease (Lee et al., 2016). On a network level, the superior frontal cortex is situated at the intersection of the default mode network, the dorsal attention network and the frontoparietal control network (Li et al., 2013; Schaefer et al., 2018; Yeo et al., 2011). This particular network embedding suggests an integrating role of superior frontal cortex connectivity to broader associative, self-reflective processes, as well as controlling operations across the cortex (Andrews-Hanna et al., 2014; Li et al., 2013; Spreng et al., 2013). Our observation of an inter-relationship of cognition and affect in superior frontal cortex is further in line with a meta-analysis showing that interactions between emotion and cognition were associated with this region, next to medial prefrontal cortex and basal ganglia (Cromheeke and Mueller, 2014). In addition, the results we obtained from our functional decoding analysis are in line with the variety of cognitive and emotional functions that previous studies have allocated to this region, such as activation related to social cognition, emotional valence and action inhibition (Bzdok et al., 2012; Cromheeke and Mueller, 2014; Hung et al., 2018). Our results thus extend previous evidence of cognitive and affective behavior integration in superior frontal cortex by showing a macrostructural overlap of cognition and affect in superior frontal cortex that is based on shared genetic effects.

### 4.3. Dissociations of affect and cognition in brain structure

Both individual differences in cognitive and affective traits could be linked to local brain structure. Total cognition was associated with lower thickness in frontal regions which is in line with some studies (Goh et al., 2011; Salat et al., 2002; Sowell et al., 2001; Van Petten et al., 2004), but contradicting others (Fjell et al., 2006; Karama et al., 2009; Narr et al., 2006). At the same time, there is some congruency with previous studies involving the location of regions critical for cognition, including mostly frontal and parietal regions (Jung and Haier, 2007), but also anterior and posterior temporal, and occipital regions (Goh et al., 2011; Menary et al., 2013). Notably, we also observed associations between cognition and various regions within the insular cortices, functionally implicated in both cognitive, but also emotional processes (Kelly et al., 2012; Lindquist et al., 2012). In addition, we found wide-spread associations in temporal, frontal and occipital lobes between local surface area and cognitive ability, but not with affective traits. Mean affect was, however, phenotypically correlated with cortical thickness in left superior frontal cortex, left lateral occipital cortex and bilateral caudate volume. It is noteworthy that affective traits showed less strong and wide-spread associations to local thickness relative to cognitive scores. This observation is in line with reports suggesting inter-regional interactions, rather than local anatomy, may encode emotional experience (Kragel and LaBar, 2016; Langner et al., 2018; Pessoa, 2008). In fact, meta-analyses of functional neuroimaging studies did not find evidence for independent brain systems that specifically relate to positive and negative valence (Lindquist et al., 2016, 2012). This suggests that the neural representation of affect is characterized by dynamic interactions between brain regions and networks rather than functional specializations of distinct locations in the brain (Kragel and LaBar, 2016; Langner et al., 2018; Pessoa, 2008). Moreover, dissociable patterns of cortical thickness and surface area in relation to behavior might also underlie genetic influences. As such, individual variation in surface area has been associated with genes expressed pre-birth, whereas cortical thickness has been related to adult-specific gene expression and emerging genetic associations with cognitive abilities throughout development (Brouwer et al., 2014; Grasby et al., 2020; Panizzon et al., 2009).

### 4.4. Limitations and conclusions

We observed converging evidence for a shared biological basis of inter-individual differences in cognition and affect combining multi-level analysis within the HCP dataset and ad-hoc meta-analytical functional decoding. At the same time, we observed that correlations within each domain were generally stronger than between cognition and affect. Further research might benefit from studying task-based, as well as physiological measures of cognitive and affective inter-individual variation to further evaluate the dynamic relation between cognitive aptitude and habitual and transient affective experience. The singular nature of the twin-based HCP sample warrants the acquisition of comparable high-resolution neuroimaging datasets including deeply phenotyped twins and families to test replication of results. Greater insight into the association between affect and cognition may be garnered by inspecting different samples, integrating more fine-grained genetic approaches with various indices of cortical anatomy. However, associations observed here were weak, and it is of note that the combination of behavioral assessments and its association with brain structure has been recently challenged. For example, Kharabian Masouleh *et al*. showed in an extensive study that the association of psychological traits and brain structure is rarely statistically significant or even reproducible in independent samples (Kharabian Masouleh et al., 2019). Additionally, Hedge and colleagues pointed out, that commonly used measurements of behavior may not be optimal to determine underlying neural correlates, due to low between-participant variability within established paradigms (Hedge et al., 2018). Here we utilized different levels of analysis to capture the association between affect and cognition. Follow-up work on the biological basis of complex behaviors may take a similar approach and integrate behavioral assessments with neuroimaging, behavioral genetics, and functional decoding. To conclude, the current work provides evidence at three levels of enquiry that cognitive abilities and affective traits are linked to partially overlapping neurobiological processes. We anticipate that the increased availability of open datasets with rich phenotyping will enable to outline more specific biological mechanisms that help describe the relationship between thoughts and feelings.

## Supporting information

Kraljevic.Schaare.2020.supplement_v2

## Acknowledgements

We would like to sincerely thank all participants and contributors to the data provided by the Human Connectome Project, WU-Minn Consortium (Principal Investigators: David Van Essen and Kamil Ugurbil; 1U54MH091657) funded by the 16 NIH Institutes and Centers that support the NIH Blueprint for Neuroscience Research; and by the McDonnell Center for Systems Neuroscience at Washington University.

This study was supported by the European Union’s Horizon 2020 Research and Innovation Programme under Grant Agreement no. 945539 (HBP SGA3). B.T.T.Y. is funded by the Singapore National Research Foundation (NRF) Fellowship (Class of 2017) and the National University of Singapore Yong Loo Lin School of Medicine (NUHSRO/2020/124/TMR/LOA). S.L.V. was supported by the Max Planck Gesellschaft (Otto Hahn award).

