## Supplementary material for "Behavioral, Anatomical and Genetic Convergence of Affect and Cognition in Superior Frontal Cortex": Kraljevic.Schaare.2020.supplement_v2

Nevena Kraljevic*[1,2], H. Lina Schaare*[1,7], Simon B. Eickhoff[1,2], Peter Kochunov[3], B. T. Thomas Yeo[4,5,6], Shahrzad Kharabian Masouleh[1,2], Sofie L. Valk[1,2,7]

[1)] *Institute of Neuroscience and Medicine (INM-7: Brain and Behaviour), Research Centre Jülich, Jülich, Germany*
[2)] *Institute of Systems Neuroscience, Heinrich Heine University Düsseldorf, Düsseldorf, Germany*
[3)] *Maryland Psychiatric Research Center, University of Maryland School of Medicine, Baltimore, Maryland, USA*
[4)] *Department of Electrical and Computer Engineering, Centre for Sleep and Cognition, Centre for Translational MR Research, N.1 Institute for Health and Institute for Digital Medicine, National University of Singapore, Singapore, Singapore*
[5)] *Athinoula A. Martinos Center for Biomedical Imaging, Massachusetts General Hospital, Charlestown, MA, USA*
[6)] *NUS Graduate School for Integrative Sciences and Engineering, National University of Singapore, Singapore, Singapore*
[7)] *Otto Hahn group Cognitive Neurogenetics, Max Planck Institute for Human Cognitive and Brain Sciences, Leipzig, Germany*

* authors contributed equally

*Corresponding authors:*

schaare[at]cbs.mpg.de; s.valk[at]fz-juelich.de

#### SUPPLEMENTARY RESULTS

*Phenotypic association of sub-scores of cognition and affect with local brain anatomy*

We evaluated the phenotypic correlation between cognition and local cortical thickness, while controlling for global thickness. We observed that cognitive sub-scores for fluid and crystallized cognition showed similar patterns of positive and negative relations to total cognition with cortical thickness (Supplementary Figure 2, Supplementary Tables 2-3). Fluid cognition was negatively associated with primarily frontal regions and positively associated with medial occipital cortex. Crystallized cognition was related to wide-spread effects in cortical thickness of frontal and parietal regions (negative associations), as well as temporal, sensorimotor and insular thickness (positive associations).

Fluid cognition was positively related to surface area primarily in bilateral occipital and temporal pole regions (Supplementary Figure 3) and crystallized cognition was positively related to surface area in bilateral inferior temporal areas, lateral frontal and parietal areas, as well as left anterior cingulate cortex (Supplementary Figure 3). See further Supplementary Tables 8-9.

With regards to subcortical volumes, we found that crystallized cognition was positively associated with bilateral hippocampal and right amygdalar volume and fluid cognition was negatively associated with left pallidum volume (Supplementary Tables 11-12).

Sub-score analyses of affect yielded that positive affect was negatively associated with cortical thickness in left superior frontal cortex, whereas negative affect was negatively associated with occipital cortical thickness (Supplementary Figure 2, Supplementary Tables 5-6). Furthermore, there were significant phenotypic associations for both positive and negative affect with bilateral caudate volumes (Supplementary Tables 14-15). Effects with surface area were not significant.

*Genetic correlation of sub-scores of cognition and affect with local brain anatomy*

To assess if the correlation between cognitive and affective traits on the one hand and local brain structure on the other is driven by shared genetic effects, genetic correlation analyses were performed through a bivariate polygenetic analysis. In general, there was a strong overlap between phenotypic correlations and genetic correlations (Supplementary Figure 5, Supplementary Figure 6). 18 out of 21 phenotypic correlations between fluid cognition, and 36 out of 42 phenotypic correlations for crystallized cognition and local cortical thickness could be attributed to genetic effects (Supplementary Table 18).

Regarding associations with surface area, we found 11 out of 14 related to fluid cognition and 36 out of 50 related to crystallized cognition to be attributable to shared genetic effects (Supplementary Table 19).

Fluid cognition was also genetically correlated with volume in the left pallidum ($\rho_g$=-0.230, p=0.003) and there was a genetic correlation between crystallized cognition and bilateral hippocampal volume (left: $\rho_g$=0.159, p=0.004; right: $\rho_g$=0.098, p=0.026; Supplementary Table 20).

Genetic correlations with affective sub-scores yielded that only the phenotypic association of positive affect with superior frontal cortex was driven by shared genetic effects (Supplementary Figure 5, Supplementary Table 18). We also found that the associations of positive, as well as negative affect with bilateral caudate volumes were genetically driven (Supplementary Tables 20).

### SUPPLEMENTARY FIGURES

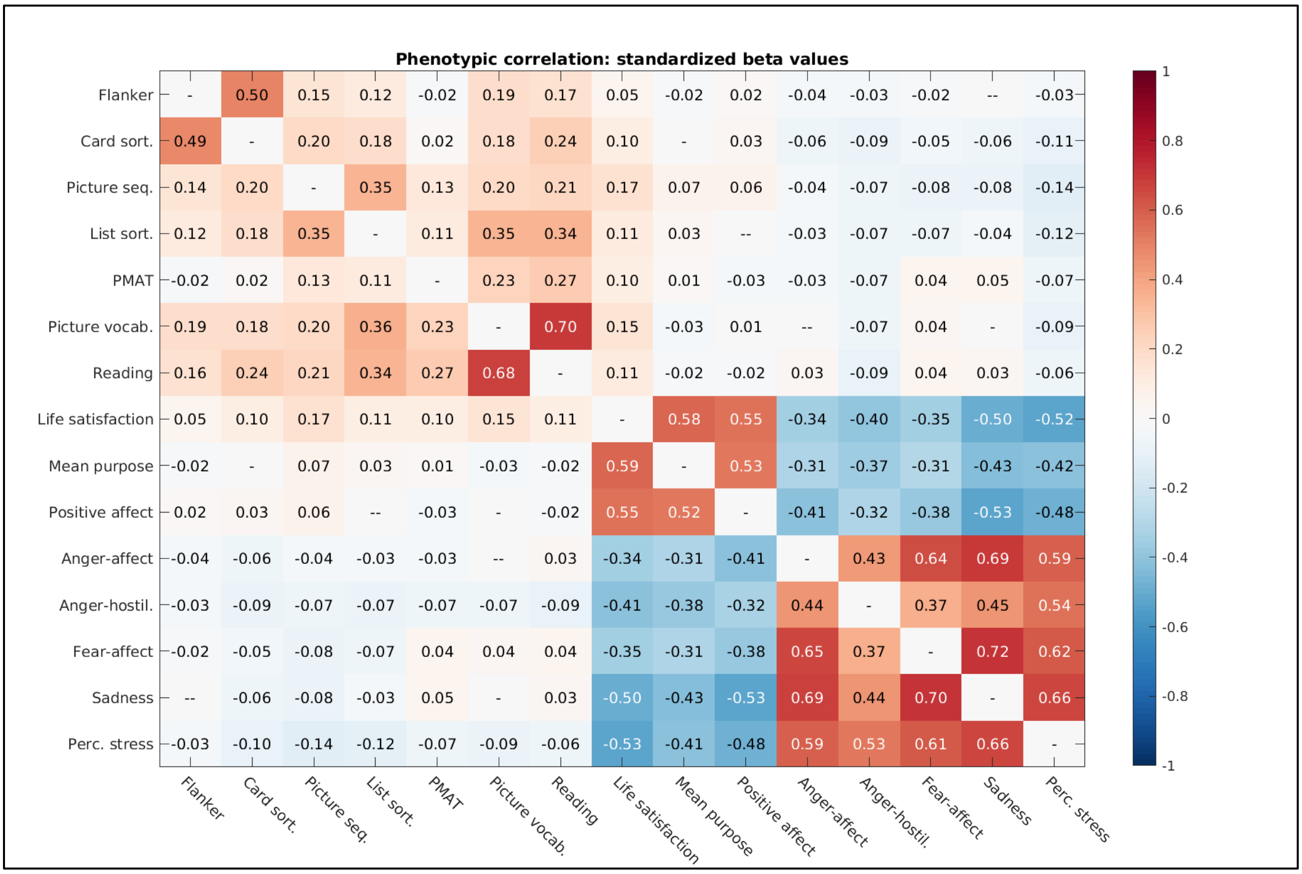

***Supplementary Figure 1. Phenotypic correlations between cognitive and affective sub-scores.*** Abbreviations: Card sort.: Dimensional Change Card Sorting, Picture seq.: Picture Sequence Memory, List sort.: List Sorting, PMAT: Penn Matrix Test pattern comparison test, Picture vocab.: Picture Vocabulary, Anger-hostil.: Anger sub-scale Hostility, Perc. stress: Perceived stress.

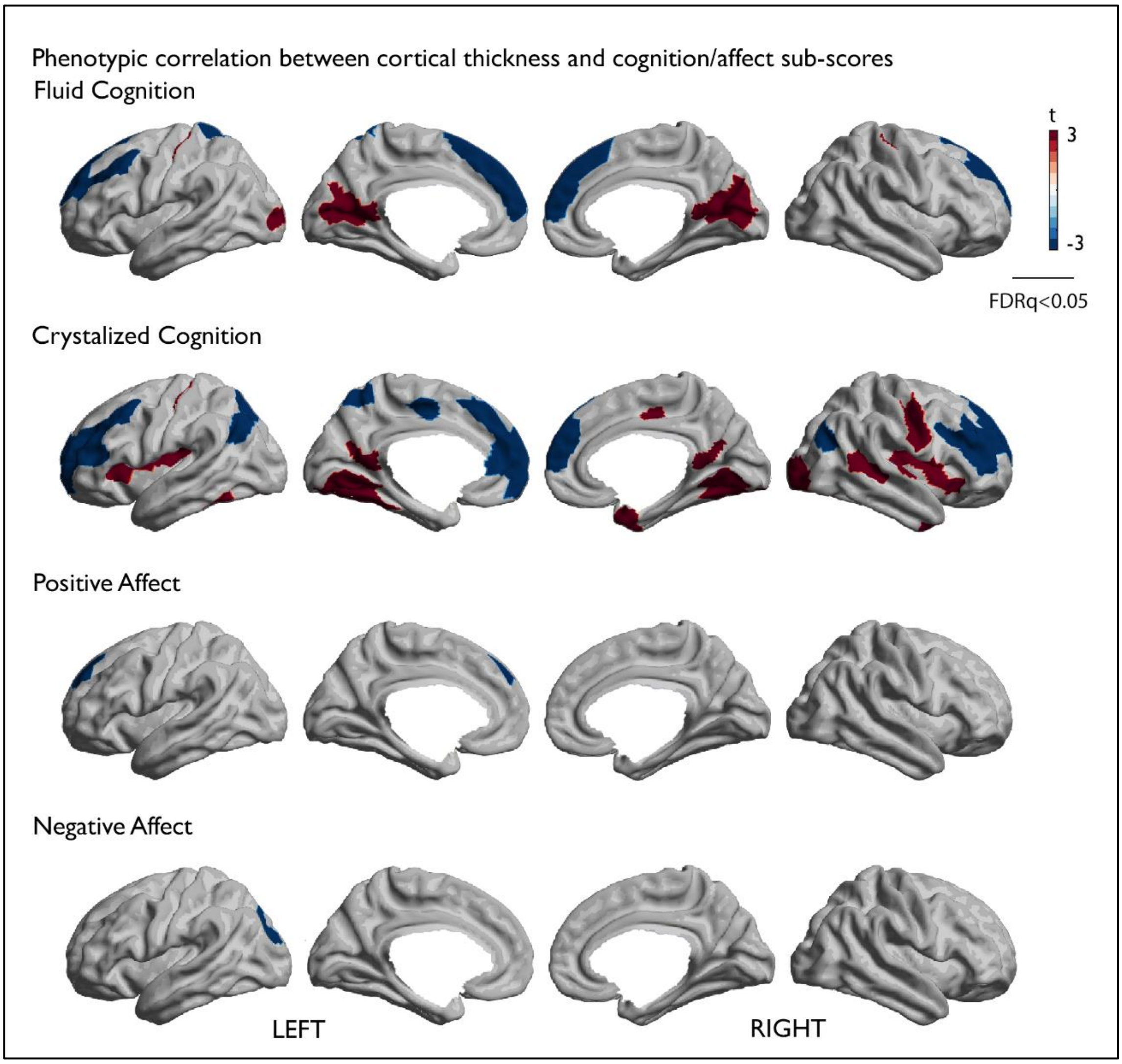

***Supplementary Figure 2. Whole-brain phenotypic correlation of cortical thickness with cognitive and affective sub-scores.*** Positive correlation is depicted in red, negative in blue. Only correlations at FDRq<0.05 are depicted

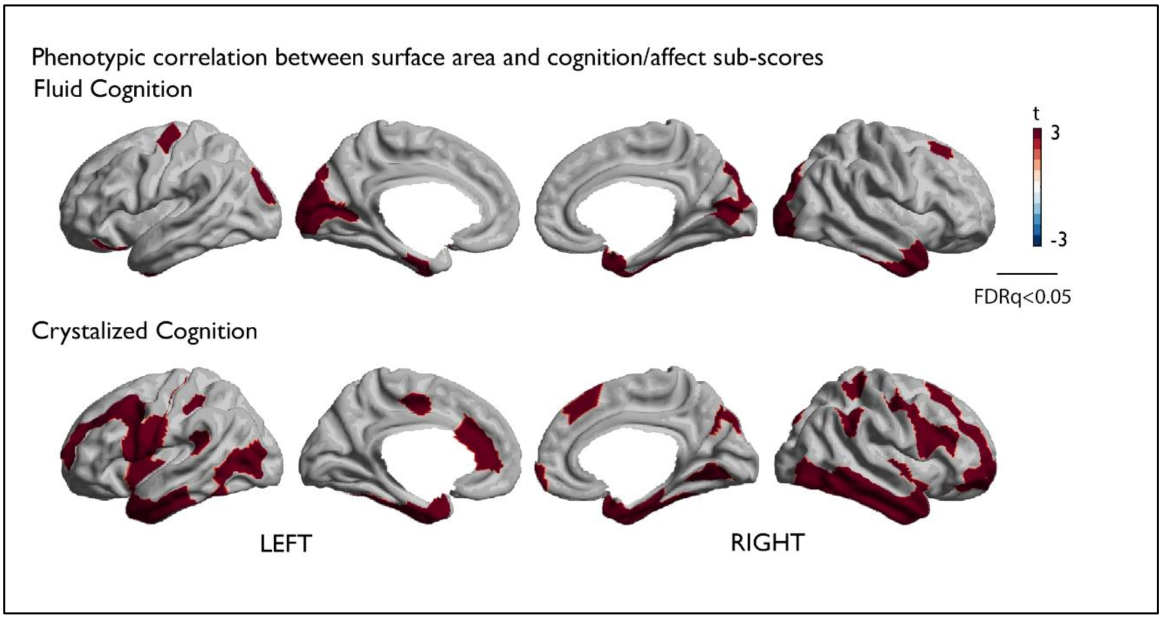

***Supplementary Figure 3. Whole-brain phenotypic correlation between surface area and cognitive sub-scores.*** Positive correlation is depicted in red, negative in blue. Only correlations at FDRq<0.05 are depicted. Associations with affective sub-scores were not significant.

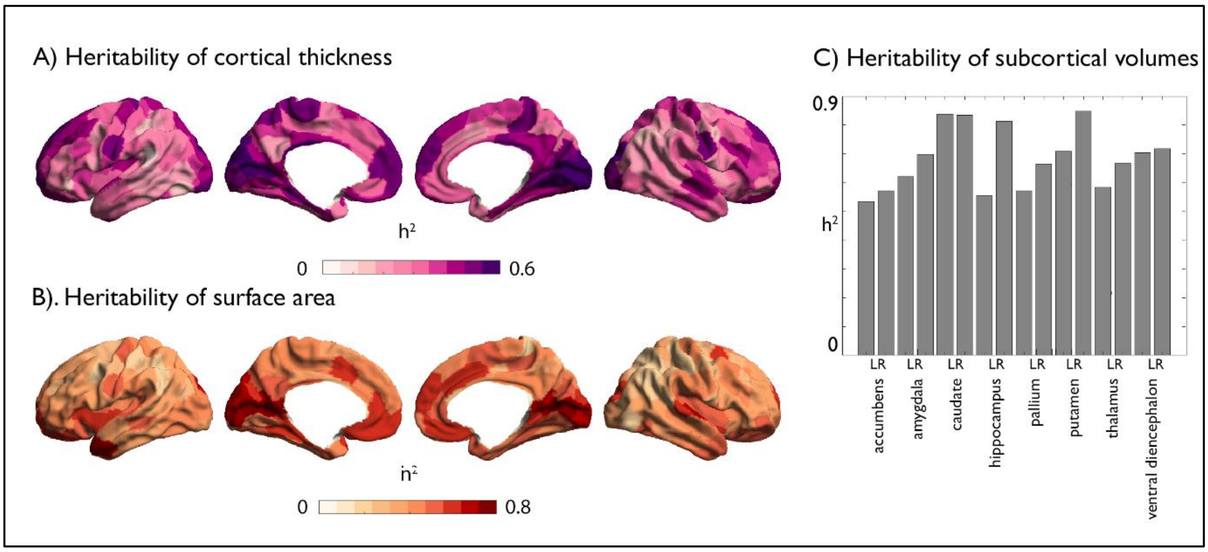

***Supplementary Figure 4. Heritability of local cortical thickness, surface area and subcortical volumes.*** Heritability of local cortical thickness, surface area and subcortical volumes. A) Heritability of local cortical thickness per parcel (200 parcel solution Schaefer, 2018); B) Heritability of local surface area per parcel. C) Heritability of subcortical volumes per FreeSurfer-segmented region.

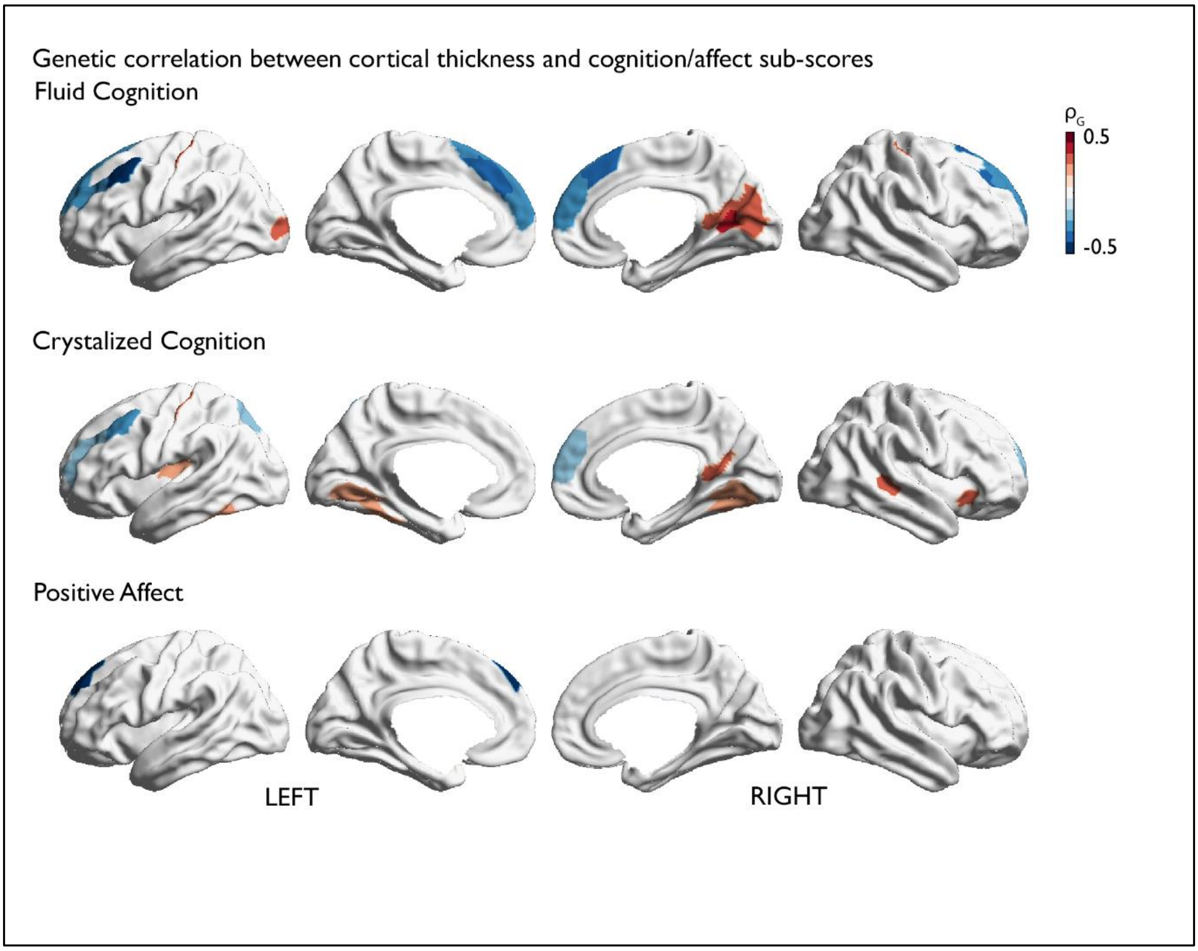

***Supplementary Figure 5. Whole-brain genetic correlation of cortical thickness with cognitive and affective sub-scores.*** Positive correlation is depicted in red, negative in blue. Only correlations at FDRq<0.05 are depicted.

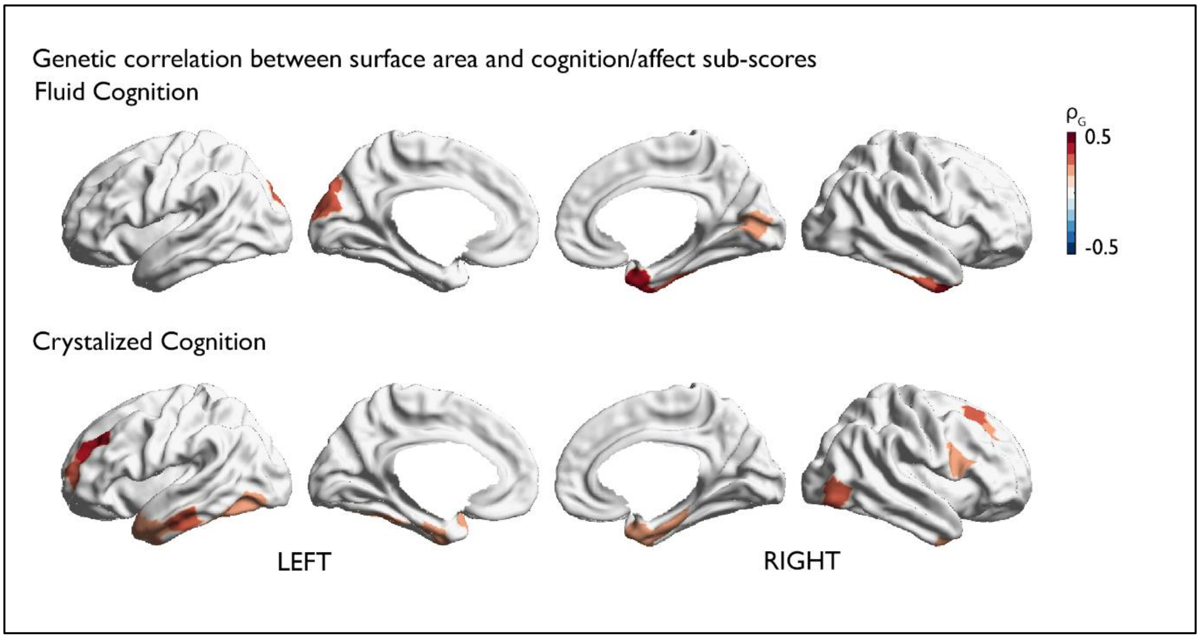

***Supplementary Figure 6. Whole-brain genetic correlation between surface area and cognitive sub-scores.*** Positive correlation is depicted in red, negative in blue. Only correlations at FDRq<0.05 are depicted. Associations with affective sub-scores were not significant.

#### SUPPLEMENTARY TABLES

***Supplementary Table 1.*** **Total cognition and local cortical thickness.**

| | Total Cognition | p | Fluid | p | Crystallized | p | Positive Affect | p | Negative Affect | p | Mean Affect | p |
|---|---|---|---|---|---|---|---|---|---|---|---|---|
| **7Networks_LH_Vis_1** | 3.634 | 0.000 | 1.637 | 0.051 | 5.041 | 0.000 | -0.192 | 0.424 | 1.351 | 0.089 | -0.842 | 0.200 |
| **7Networks_LH_Vis_4** | 3.849 | 0.000 | 2.505 | 0.006 | 4.220 | 0.000 | 0.341 | 0.367 | -0.648 | 0.258 | 0.547 | 0.292 |
| **7Networks_LH_Vis_9** | 2.905 | 0.002 | 3.161 | 0.001 | 1.806 | 0.036 | 2.439 | 0.007 | -1.932 | 0.027 | 2.451 | 0.007 |
| **7Networks_LH_Vis_10** | 3.245 | 0.001 | 3.542 | 0.000 | 1.765 | 0.039 | 0.195 | 0.423 | -0.858 | 0.196 | 0.577 | 0.282 |
| **7Networks_LH_SomMot_3** | 3.962 | 0.000 | 2.528 | 0.006 | 4.312 | 0.000 | 0.425 | 0.336 | -1.248 | 0.106 | 0.921 | 0.179 |
| **7Networks_LH_SomMot_10** | 5.528 | 0.000 | 4.323 | 0.000 | 5.358 | 0.000 | 2.074 | 0.019 | -1.915 | 0.028 | 2.232 | 0.013 |
| **7Networks_LH_DorsAttn_Post_7** | -2.965 | 0.002 | -1.553 | 0.060 | -3.896 | 0.000 | -2.011 | 0.022 | 0.779 | 0.218 | -1.579 | 0.057 |
| **7Networks_LH_DorsAttn_Post_10** | -3.090 | 0.001 | -3.113 | 0.001 | -2.287 | 0.011 | -0.634 | 0.263 | -0.256 | 0.399 | -0.225 | 0.411 |
| **7Networks_LH_DorsAttn_FEF_2** | -3.016 | 0.001 | -2.613 | 0.005 | -2.653 | 0.004 | -2.759 | 0.003 | 2.216 | 0.013 | -2.791 | 0.003 |
| **7Networks_LH_SalVentAttn_FrOperIns_2** | 3.597 | 0.000 | 2.826 | 0.002 | 3.433 | 0.000 | 0.578 | 0.282 | 0.694 | 0.244 | -0.044 | 0.483 |
| **7Networks_LH_SalVentAttn_PFCl_1** | -3.875 | 0.000 | -3.047 | 0.001 | -3.744 | 0.000 | -0.433 | 0.333 | 0.062 | 0.475 | -0.282 | 0.389 |
| **7Networks_LH_Default_Temp_5** | 2.995 | 0.001 | 2.368 | 0.009 | 2.399 | 0.008 | 0.564 | 0.286 | -1.098 | 0.136 | 0.920 | 0.179 |
| **7Networks_LH_Default_PFC_4** | -3.361 | 0.000 | -2.920 | 0.002 | -2.830 | 0.002 | -2.146 | 0.016 | 2.116 | 0.017 | -2.383 | 0.009 |
| **7Networks_LH_Default_PFC_5** | -2.891 | 0.002 | -2.182 | 0.015 | -2.729 | 0.003 | -2.378 | 0.009 | 2.351 | 0.009 | -2.644 | 0.004 |
| **7Networks_LH_Default_PFC_7** | -4.608 | 0.000 | -3.923 | 0.000 | -4.138 | 0.000 | -1.869 | 0.031 | 1.460 | 0.072 | -1.867 | 0.031 |
| **7Networks_LH_Default_PFC_9** | -3.236 | 0.001 | -3.203 | 0.001 | -2.227 | 0.013 | -3.972 | 0.000 | 3.257 | 0.001 | -4.056 | 0.000 |
| **7Networks_LH_Default_PFC_10** | -4.396 | 0.000 | -4.220 | 0.000 | -3.436 | 0.000 | -1.728 | 0.042 | 1.211 | 0.113 | -1.651 | 0.050 |
| **7Networks_LH_Default_PFC_11** | -6.733 | 0.000 | -5.354 | 0.000 | -6.197 | 0.000 | -2.349 | 0.010 | 1.307 | 0.096 | -2.060 | 0.020 |
| **7Networks_LH_Default_PFC_13** | -3.544 | 0.000 | -3.644 | 0.000 | -2.594 | 0.005 | -3.002 | 0.001 | 2.874 | 0.002 | -3.288 | 0.001 |
| **7Networks_LH_Default_pCunPCC_1** | 3.841 | 0.000 | 3.059 | 0.001 | 3.528 | 0.000 | 0.482 | 0.315 | 0.097 | 0.461 | 0.225 | 0.411 |
| **7Networks_RH_Vis_4** | 3.017 | 0.001 | 0.599 | 0.275 | 5.039 | 0.000 | 0.770 | 0.221 | 0.251 | 0.401 | 0.307 | 0.379 |
| **7Networks_RH_Vis_9** | 2.945 | 0.002 | 3.524 | 0.000 | 1.345 | 0.089 | -0.816 | 0.207 | 0.613 | 0.270 | -0.802 | 0.211 |
| **7Networks_RH_Vis_10** | 3.274 | 0.001 | 3.447 | 0.000 | 1.819 | 0.035 | 1.610 | 0.054 | -1.868 | 0.031 | 1.940 | 0.026 |

| | | | | | | | | | | | | |
|---|---|---|---|---|---|---|---|---|---|---|---|---|
| **7Networks_RH_Vis_13** | 3.559 | 0.000 | 3.323 | 0.000 | 2.686 | 0.004 | -0.244 | 0.404 | -0.665 | 0.253 | 0.220 | 0.413 |
| **7Networks_RH_SomMot_1** | 3.364 | 0.000 | 2.182 | 0.015 | 3.694 | 0.000 | 0.480 | 0.316 | -0.429 | 0.334 | 0.509 | 0.305 |
| **7Networks_RH_SomMot_7** | 3.115 | 0.001 | 2.183 | 0.015 | 3.026 | 0.001 | -0.129 | 0.448 | 0.261 | 0.397 | -0.216 | 0.415 |
| **7Networks_RH_SomMot_8** | 3.432 | 0.000 | 2.027 | 0.021 | 3.730 | 0.000 | 2.468 | 0.007 | -1.880 | 0.030 | 2.440 | 0.007 |
| **7Networks_RH_SomMot_12** | 3.623 | 0.000 | 3.565 | 0.000 | 2.463 | 0.007 | 2.166 | 0.015 | -1.347 | 0.089 | 1.977 | 0.024 |
| **7Networks_RH_Cont_PFCv_1** | 3.128 | 0.001 | 2.212 | 0.014 | 3.075 | 0.001 | 2.337 | 0.010 | -0.089 | 0.464 | 1.391 | 0.082 |
| **7Networks_RH_Cont_PFCl_5** | -3.145 | 0.001 | -2.111 | 0.018 | -3.348 | 0.000 | -1.088 | 0.138 | 1.322 | 0.093 | -1.343 | 0.090 |
| **7Networks_RH_Cont_PFCl_6** | -3.404 | 0.000 | -2.021 | 0.022 | -4.262 | 0.000 | -2.152 | 0.016 | 1.456 | 0.073 | -2.028 | 0.021 |
| **7Networks_RH_Cont_PFCl_7** | -2.892 | 0.002 | -2.327 | 0.010 | -2.423 | 0.008 | -2.029 | 0.021 | 1.750 | 0.040 | -2.117 | 0.017 |
| **7Networks_RH_Cont_PFCmp_2** | -3.351 | 0.000 | -3.066 | 0.001 | -2.724 | 0.003 | -0.906 | 0.183 | 2.161 | 0.015 | -1.693 | 0.045 |
| **7Networks_RH_Default_PFCdPFCm_4** | -4.405 | 0.000 | -3.151 | 0.001 | -4.669 | 0.000 | -2.079 | 0.019 | 1.819 | 0.035 | -2.183 | 0.015 |
| **7Networks_RH_Default_PFCdPFCm_5** | -5.419 | 0.000 | -5.028 | 0.000 | -4.067 | 0.000 | -2.601 | 0.005 | 3.632 | 0.000 | -3.468 | 0.000 |
| **7Networks_RH_Default_PFCdPFCm_6** | -3.590 | 0.000 | -3.382 | 0.000 | -2.756 | 0.003 | -2.126 | 0.017 | 1.594 | 0.056 | -2.088 | 0.019 |
| **7Networks_RH_Default_pCunPCC_1** | 5.853 | 0.000 | 4.358 | 0.000 | 5.813 | 0.000 | 1.263 | 0.103 | -1.571 | 0.058 | 1.578 | 0.057 |

***Supplementary Table 2.*** **Fluid cognition and local cortical thickness.**

| | **Total Cognition** | **p** | **Fluid** | **p** | **Crystallized** | **p** | **Positive Affect** | **p** | **Negative Affect** | **p** | **Mean Affect** | **p** |
|---|---|---|---|---|---|---|---|---|---|---|---|---|
| **7Networks_LH_Vis_9** | 2.905 | 0.002 | 3.161 | 0.001 | 1.806 | 0.036 | 2.439 | 0.007 | -1.932 | 0.027 | 2.451 | 0.007 |
| **7Networks_LH_Vis_10** | 3.245 | 0.001 | 3.542 | 0.000 | 1.765 | 0.039 | 0.195 | 0.423 | -0.858 | 0.196 | 0.577 | 0.282 |
| **7Networks_LH_Vis_12** | 2.282 | 0.011 | 3.201 | 0.001 | 0.349 | 0.363 | -0.064 | 0.474 | -2.275 | 0.012 | 1.194 | 0.116 |
| **7Networks_LH_SomMot_10** | 5.528 | 0.000 | 4.323 | 0.000 | 5.358 | 0.000 | 2.074 | 0.019 | -1.915 | 0.028 | 2.232 | 0.013 |
| **7Networks_LH_DorsAttn_Post_10** | -3.090 | 0.001 | -3.113 | 0.001 | -2.287 | 0.011 | -0.634 | 0.263 | -0.256 | 0.399 | -0.225 | 0.411 |
| **7Networks_LH_SalVentAttn_PFCl_1** | -3.875 | 0.000 | -3.047 | 0.001 | -3.744 | 0.000 | -0.433 | 0.333 | 0.062 | 0.475 | -0.282 | 0.389 |
| **7Networks_LH_Default_PFC_7** | -4.608 | 0.000 | -3.923 | 0.000 | -4.138 | 0.000 | -1.869 | 0.031 | 1.460 | 0.072 | -1.867 | 0.031 |
| **7Networks_LH_Default_PFC_9** | -3.236 | 0.001 | -3.203 | 0.001 | -2.227 | 0.013 | -3.972 | 0.000 | 3.257 | 0.001 | -4.056 | 0.000 |
| **7Networks_LH_Default_PFC_10** | -4.396 | 0.000 | -4.220 | 0.000 | -3.436 | 0.000 | -1.728 | 0.042 | 1.211 | 0.113 | -1.651 | 0.050 |
| **7Networks_LH_Default_PFC_11** | -6.733 | 0.000 | -5.354 | 0.000 | -6.197 | 0.000 | -2.349 | 0.010 | 1.307 | 0.096 | -2.060 | 0.020 |

| | | | | | | | | | | | | |
|---|---|---|---|---|---|---|---|---|---|---|---|---|
| **7Networks_LH_Default_PFC_13** | -3.544 | 0.000 | -3.644 | 0.000 | -2.594 | 0.005 | -3.002 | 0.001 | 2.874 | 0.002 | -3.288 | 0.001 |
| **7Networks_LH_Default_pCunPCC_1** | 3.841 | 0.000 | 3.059 | 0.001 | 3.528 | 0.000 | 0.482 | 0.315 | 0.097 | 0.461 | 0.225 | 0.411 |
| **7Networks_RH_Vis_9** | 2.945 | 0.002 | 3.524 | 0.000 | 1.345 | 0.089 | -0.816 | 0.207 | 0.613 | 0.270 | -0.802 | 0.211 |
| **7Networks_RH_Vis_10** | 3.274 | 0.001 | 3.447 | 0.000 | 1.819 | 0.035 | 1.610 | 0.054 | -1.868 | 0.031 | 1.940 | 0.026 |
| **7Networks_RH_Vis_13** | 3.559 | 0.000 | 3.323 | 0.000 | 2.686 | 0.004 | -0.244 | 0.404 | -0.665 | 0.253 | 0.220 | 0.413 |
| **7Networks_RH_SomMot_12** | 3.623 | 0.000 | 3.565 | 0.000 | 2.463 | 0.007 | 2.166 | 0.015 | -1.347 | 0.089 | 1.977 | 0.024 |
| **7Networks_RH_Cont_PFCmp_2** | -3.351 | 0.000 | -3.066 | 0.001 | -2.724 | 0.003 | -0.906 | 0.183 | 2.161 | 0.015 | -1.693 | 0.045 |
| **7Networks_RH_Default_PFCdPFCm_4** | -4.405 | 0.000 | -3.151 | 0.001 | -4.669 | 0.000 | -2.079 | 0.019 | 1.819 | 0.035 | -2.183 | 0.015 |
| **7Networks_RH_Default_PFCdPFCm_5** | -5.419 | 0.000 | -5.028 | 0.000 | -4.067 | 0.000 | -2.601 | 0.005 | 3.632 | 0.000 | -3.468 | 0.000 |
| **7Networks_RH_Default_PFCdPFCm_6** | -3.590 | 0.000 | -3.382 | 0.000 | -2.756 | 0.003 | -2.126 | 0.017 | 1.594 | 0.056 | -2.088 | 0.019 |
| **7Networks_RH_Default_pCunPCC_1** | 5.853 | 0.000 | 4.358 | 0.000 | 5.813 | 0.000 | 1.263 | 0.103 | -1.571 | 0.058 | 1.578 | 0.057 |

***Supplementary Table 3.*** **Crystallized cognition and local cortical thickness.**

| | **Total Cognition** | **p** | **Fluid** | **p** | **Crystallized** | **p** | **Positive Affect** | **p** | **Negative Affect** | **p** | **Mean Affect** | **p** |
|---|---|---|---|---|---|---|---|---|---|---|---|---|
| **7Networks_LH_Vis_1** | 3.634 | 0.000 | 1.637 | 0.051 | 5.041 | 0.000 | -0.192 | 0.424 | 1.351 | 0.089 | -0.842 | 0.200 |
| **7Networks_LH_Vis_2** | 2.500 | 0.006 | 0.692 | 0.245 | 4.025 | 0.000 | 0.066 | 0.474 | 0.594 | 0.276 | -0.284 | 0.388 |
| **7Networks_LH_Vis_4** | 3.849 | 0.000 | 2.505 | 0.006 | 4.220 | 0.000 | 0.341 | 0.367 | -0.648 | 0.258 | 0.547 | 0.292 |
| **7Networks_LH_SomMot_3** | 3.962 | 0.000 | 2.528 | 0.006 | 4.312 | 0.000 | 0.425 | 0.336 | -1.248 | 0.106 | 0.921 | 0.179 |
| **7Networks_LH_SomMot_8** | -2.063 | 0.020 | -0.909 | 0.182 | -2.996 | 0.001 | 0.562 | 0.287 | -0.356 | 0.361 | 0.517 | 0.303 |
| **7Networks_LH_SomMot_10** | 5.528 | 0.000 | 4.323 | 0.000 | 5.358 | 0.000 | 2.074 | 0.019 | -1.915 | 0.028 | 2.232 | 0.013 |
| **7Networks_LH_DorsAttn_Post_1** | 2.215 | 0.013 | 1.084 | 0.139 | 3.007 | 0.001 | 0.626 | 0.266 | -1.264 | 0.103 | 1.045 | 0.148 |
| **7Networks_LH_DorsAttn_Post_7** | -2.965 | 0.002 | -1.553 | 0.060 | -3.896 | 0.000 | -2.011 | 0.022 | 0.779 | 0.218 | -1.579 | 0.057 |
| **7Networks_LH_DorsAttn_Post_9** | -2.598 | 0.005 | -1.605 | 0.054 | -2.896 | 0.002 | 1.108 | 0.134 | -0.885 | 0.188 | 1.117 | 0.132 |
| **7Networks_LH_SalVentAttn_FrOperIns_2** | 3.597 | 0.000 | 2.826 | 0.002 | 3.433 | 0.000 | 0.578 | 0.282 | 0.694 | 0.244 | -0.044 | 0.483 |
| **7Networks_LH_SalVentAttn_FrOperIns_3** | 1.726 | 0.042 | 0.326 | 0.372 | 3.055 | 0.001 | 0.243 | 0.404 | 0.212 | 0.416 | 0.025 | 0.490 |
| **7Networks_LH_SalVentAttn_PFCl_1** | -3.875 | 0.000 | -3.047 | 0.001 | -3.744 | 0.000 | -0.433 | 0.333 | 0.062 | 0.475 | -0.282 | 0.389 |
| **7Networks_LH_Cont_PFCl_2** | -2.534 | 0.006 | -1.334 | 0.091 | -2.868 | 0.002 | -1.357 | 0.087 | 0.173 | 0.432 | -0.874 | 0.191 |

| | | | | | | | | | | | | |
|---|---|---|---|---|---|---|---|---|---|---|---|---|
| **7Networks_LH_Cont_PFCl_3** | -2.402 | 0.008 | -1.092 | 0.138 | -3.294 | 0.001 | -1.719 | 0.043 | 2.723 | 0.003 | -2.466 | 0.007 |
| **7Networks_LH_Default_Par_4** | -1.484 | 0.069 | 0.017 | 0.493 | -3.068 | 0.001 | 2.515 | 0.006 | -1.342 | 0.090 | 2.174 | 0.015 |
| **7Networks_LH_Default_PFC_4** | -3.361 | 0.000 | -2.920 | 0.002 | -2.830 | 0.002 | -2.146 | 0.016 | 2.116 | 0.017 | -2.383 | 0.009 |
| **7Networks_LH_Default_PFC_6** | -2.609 | 0.005 | -1.717 | 0.043 | -2.897 | 0.002 | -1.749 | 0.040 | -0.003 | 0.499 | -1.003 | 0.158 |
| **7Networks_LH_Default_PFC_7** | -4.608 | 0.000 | -3.923 | 0.000 | -4.138 | 0.000 | -1.869 | 0.031 | 1.460 | 0.072 | -1.867 | 0.031 |
| **7Networks_LH_Default_PFC_10** | -4.396 | 0.000 | -4.220 | 0.000 | -3.436 | 0.000 | -1.728 | 0.042 | 1.211 | 0.113 | -1.651 | 0.050 |
| **7Networks_LH_Default_PFC_11** | -6.733 | 0.000 | -5.354 | 0.000 | -6.197 | 0.000 | -2.349 | 0.010 | 1.307 | 0.096 | -2.060 | 0.020 |
| **7Networks_LH_Default_pCunPCC_1** | 3.841 | 0.000 | 3.059 | 0.001 | 3.528 | 0.000 | 0.482 | 0.315 | 0.097 | 0.461 | 0.225 | 0.411 |
| **7Networks_RH_Vis_3** | 1.023 | 0.153 | -1.123 | 0.131 | 3.535 | 0.000 | 1.247 | 0.106 | 0.641 | 0.261 | 0.369 | 0.356 |
| **7Networks_RH_Vis_4** | 3.017 | 0.001 | 0.599 | 0.275 | 5.039 | 0.000 | 0.770 | 0.221 | 0.251 | 0.401 | 0.307 | 0.379 |
| **7Networks_RH_Vis_8** | 2.370 | 0.009 | 0.972 | 0.166 | 3.390 | 0.000 | 2.309 | 0.011 | -1.035 | 0.151 | 1.888 | 0.030 |
| **7Networks_RH_SomMot_1** | 3.364 | 0.000 | 2.182 | 0.015 | 3.694 | 0.000 | 0.480 | 0.316 | -0.429 | 0.334 | 0.509 | 0.305 |
| **7Networks_RH_SomMot_3** | 2.108 | 0.018 | 1.115 | 0.132 | 3.015 | 0.001 | 1.678 | 0.047 | -1.483 | 0.069 | 1.770 | 0.039 |
| **7Networks_RH_SomMot_7** | 3.115 | 0.001 | 2.183 | 0.015 | 3.026 | 0.001 | -0.129 | 0.448 | 0.261 | 0.397 | -0.216 | 0.415 |
| **7Networks_RH_SomMot_8** | 3.432 | 0.000 | 2.027 | 0.021 | 3.730 | 0.000 | 2.468 | 0.007 | -1.880 | 0.030 | 2.440 | 0.007 |
| **7Networks_RH_SomMot_10** | 2.545 | 0.006 | 1.803 | 0.036 | 2.796 | 0.003 | -0.303 | 0.381 | -0.601 | 0.274 | 0.151 | 0.440 |
| **7Networks_RH_SalVentAttn_TempOccPar_1** | 2.674 | 0.004 | 1.473 | 0.071 | 3.147 | 0.001 | 0.866 | 0.193 | -1.108 | 0.134 | 1.099 | 0.136 |
| **7Networks_RH_SalVentAttn_FrOperIns_4** | 1.849 | 0.032 | 0.470 | 0.319 | 3.247 | 0.001 | 0.926 | 0.177 | -0.218 | 0.414 | 0.650 | 0.258 |
| **7Networks_RH_Limbic_TempPole_1** | 2.334 | 0.010 | 0.885 | 0.188 | 3.289 | 0.001 | 0.312 | 0.377 | -0.033 | 0.487 | 0.198 | 0.422 |
| **7Networks_RH_Cont_PFCv_1** | 3.128 | 0.001 | 2.212 | 0.014 | 3.075 | 0.001 | 2.337 | 0.010 | -0.089 | 0.464 | 1.391 | 0.082 |
| **7Networks_RH_Cont_PFCl_3** | -2.727 | 0.003 | -1.658 | 0.049 | -3.094 | 0.001 | 0.476 | 0.317 | 0.562 | 0.287 | -0.031 | 0.488 |
| **7Networks_RH_Cont_PFCl_4** | -2.375 | 0.009 | -1.065 | 0.144 | -3.064 | 0.001 | -1.252 | 0.105 | 1.026 | 0.152 | -1.277 | 0.101 |
| **7Networks_RH_Cont_PFCl_5** | -3.145 | 0.001 | -2.111 | 0.018 | -3.348 | 0.000 | -1.088 | 0.138 | 1.322 | 0.093 | -1.343 | 0.090 |
| **7Networks_RH_Cont_PFCl_6** | -3.404 | 0.000 | -2.021 | 0.022 | -4.262 | 0.000 | -2.152 | 0.016 | 1.456 | 0.073 | -2.028 | 0.021 |
| **7Networks_RH_Default_Par_1** | -1.667 | 0.048 | -0.223 | 0.412 | -2.852 | 0.002 | -0.023 | 0.491 | 0.424 | 0.336 | -0.243 | 0.404 |
| **7Networks_RH_Default_Temp_5** | 2.767 | 0.003 | 0.891 | 0.186 | 3.923 | 0.000 | 0.651 | 0.258 | -0.611 | 0.271 | 0.706 | 0.240 |
| **7Networks_RH_Default_PFCdPFCm_4** | -4.405 | 0.000 | -3.151 | 0.001 | -4.669 | 0.000 | -2.079 | 0.019 | 1.819 | 0.035 | -2.183 | 0.015 |
| **7Networks_RH_Default_PFCdPFCm_5** | -5.419 | 0.000 | -5.028 | 0.000 | -4.067 | 0.000 | -2.601 | 0.005 | 3.632 | 0.000 | -3.468 | 0.000 |
| **7Networks_RH_Default_pCunPCC_1** | 5.853 | 0.000 | 4.358 | 0.000 | 5.813 | 0.000 | 1.263 | 0.103 | -1.571 | 0.058 | 1.578 | 0.057 |

*Supplementary Table 4.* **Mean affect and local cortical thickness.**

| | Total Cognition | p | Fluid | p | Crystallized | p | Positive Affect | p | Negative Affect | p | Mean Affect | p |
|---|---|---|---|---|---|---|---|---|---|---|---|---|
| **7Networks_LH_Vis_14** | -0.299 | 0.382 | -0.098 | 0.461 | -0.491 | 0.312 | 2.950 | 0.002 | -4.241 | 0.000 | 4.000 | 0.000 |
| **7Networks_LH_Default_PFC_9** | -3.236 | 0.001 | -3.203 | 0.001 | -2.227 | 0.013 | -3.972 | 0.000 | 3.257 | 0.001 | -4.056 | 0.000 |
| **7Networks_RH_DorsAttn_Post_5** | 0.538 | 0.295 | 1.056 | 0.146 | -0.536 | 0.296 | 3.274 | 0.001 | -2.974 | 0.002 | 3.499 | 0.000 |

*Supplementary Table 5.* **Positive affect and local cortical thickness.**

| | Total Cognition | p | Fluid | p | Crystallized | p | Positive Affect | p | Negative Affect | p | Mean Affect | p |
|---|---|---|---|---|---|---|---|---|---|---|---|---|
| **7Networks_LH_Default_PFC_9** | -3.236 | 0.001 | -3.203 | 0.001 | -2.227 | 0.013 | -3.972 | 0.000 | 3.257 | 0.001 | -4.056 | 0.000 |

*Supplementary Table 6.* **Negative affect and local cortical thickness.**

| | Total Cognition | p | Fluid | p | Crystallized | p | Positive Affect | p | Negative Affect | p | Mean Affect | p |
|---|---|---|---|---|---|---|---|---|---|---|---|---|
| **7Networks_LH_Vis_14** | -0.299 | 0.382 | -0.098 | 0.461 | -0.491 | 0.312 | 2.950 | 0.002 | -4.241 | 0.000 | 4.000 | 0.000 |

*Supplementary Table 7.* **Total cognition and local surface area.**

| | Total Cognition | p | Fluid | p | Crystallized | p | Positive Affect | p | Negative Affect | p | Mean Affect | p |
|---|---|---|---|---|---|---|---|---|---|---|---|---|
| **7Networks_LH_Vis_7** | 2.743 | 0.003 | 3.091 | 0.001 | 1.092 | 0.138 | 0.102 | 0.459 | -1.493 | 0.068 | 0.869 | 0.192 |
| **7Networks_LH_Vis_9** | 2.578 | 0.005 | 2.602 | 0.005 | 0.973 | 0.165 | 0.274 | 0.392 | -1.406 | 0.080 | 0.921 | 0.179 |
| **7Networks_LH_Vis_10** | 2.665 | 0.004 | 2.942 | 0.002 | 1.187 | 0.118 | 0.848 | 0.198 | -1.311 | 0.095 | 1.200 | 0.115 |
| **7Networks_LH_Vis_11** | 2.755 | 0.003 | 1.996 | 0.023 | 2.568 | 0.005 | 0.610 | 0.271 | -1.889 | 0.030 | 1.376 | 0.085 |
| **7Networks_LH_Vis_13** | 4.304 | 0.000 | 4.344 | 0.000 | 2.332 | 0.010 | 1.986 | 0.024 | -1.734 | 0.042 | 2.083 | 0.019 |
| **7Networks_LH_Vis_14** | 2.863 | 0.002 | 2.994 | 0.001 | 1.520 | 0.064 | 1.976 | 0.024 | -0.566 | 0.286 | 1.441 | 0.075 |

| | | | | | | | | | | | | |
|---|---|---|---|---|---|---|---|---|---|---|---|---|
| **7Networks_LH_SomMot_8** | 3.059 | 0.001 | 2.224 | 0.013 | 2.947 | 0.002 | 1.751 | 0.040 | -1.605 | 0.054 | 1.878 | 0.030 |
| **7Networks_LH_SomMot_10** | 3.065 | 0.001 | 2.251 | 0.012 | 3.006 | 0.001 | -0.806 | 0.210 | 0.462 | 0.322 | -0.714 | 0.238 |
| **7Networks_LH_SomMot_12** | 3.113 | 0.001 | 2.916 | 0.002 | 2.259 | 0.012 | -0.318 | 0.375 | -0.357 | 0.360 | 0.011 | 0.495 |
| **7Networks_LH_SalVentAttn_Med_2** | 2.860 | 0.002 | 2.884 | 0.002 | 1.764 | 0.039 | 1.059 | 0.145 | -0.603 | 0.273 | 0.936 | 0.175 |
| **7Networks_LH_Limbic_OFC_1** | 3.075 | 0.001 | 3.644 | 0.000 | 1.421 | 0.078 | 1.515 | 0.065 | -1.077 | 0.141 | 1.455 | 0.073 |
| **7Networks_LH_Limbic_TempPole_1** | 4.959 | 0.000 | 3.330 | 0.000 | 4.951 | 0.000 | 1.306 | 0.096 | -1.627 | 0.052 | 1.634 | 0.051 |
| **7Networks_LH_Limbic_TempPole_2** | 3.216 | 0.001 | 2.079 | 0.019 | 3.297 | 0.001 | 1.862 | 0.031 | -1.545 | 0.061 | 1.909 | 0.028 |
| **7Networks_LH_Limbic_TempPole_3** | 4.238 | 0.000 | 2.830 | 0.002 | 4.458 | 0.000 | 1.039 | 0.149 | -0.603 | 0.273 | 0.924 | 0.178 |
| **7Networks_LH_Limbic_TempPole_4** | 3.346 | 0.000 | 2.135 | 0.017 | 3.898 | 0.000 | 0.620 | 0.268 | -0.210 | 0.417 | 0.470 | 0.319 |
| **7Networks_LH_Cont_PFCl_2** | 3.517 | 0.000 | 1.844 | 0.033 | 4.184 | 0.000 | 0.917 | 0.180 | -0.823 | 0.205 | 0.974 | 0.165 |
| **7Networks_LH_Cont_PFCl_5** | 3.376 | 0.000 | 1.633 | 0.051 | 4.074 | 0.000 | 0.224 | 0.411 | 1.590 | 0.056 | -0.734 | 0.232 |
| **7Networks_LH_Default_Temp_1** | 3.847 | 0.000 | 2.058 | 0.020 | 4.426 | 0.000 | 1.469 | 0.071 | -1.016 | 0.155 | 1.396 | 0.082 |
| **7Networks_LH_Default_Temp_2** | 4.388 | 0.000 | 2.659 | 0.004 | 4.597 | 0.000 | 1.333 | 0.091 | -1.205 | 0.114 | 1.420 | 0.078 |
| **7Networks_LH_Default_PFC_1** | 3.091 | 0.001 | 2.739 | 0.003 | 2.299 | 0.011 | 1.831 | 0.034 | -1.831 | 0.034 | 2.047 | 0.020 |
| **7Networks_LH_Default_PFC_6** | 3.367 | 0.000 | 2.363 | 0.009 | 3.450 | 0.000 | 0.006 | 0.498 | -0.641 | 0.261 | 0.351 | 0.363 |
| **7Networks_LH_Default_PFC_8** | 2.630 | 0.004 | 1.816 | 0.035 | 2.689 | 0.004 | 0.018 | 0.493 | -0.626 | 0.266 | 0.350 | 0.363 |
| **7Networks_LH_Default_PFC_11** | 2.827 | 0.002 | 1.209 | 0.113 | 3.639 | 0.000 | 0.608 | 0.272 | 0.802 | 0.211 | -0.087 | 0.466 |
| **7Networks_RH_Vis_4** | 3.155 | 0.001 | 2.349 | 0.009 | 2.774 | 0.003 | 0.991 | 0.161 | -0.460 | 0.323 | 0.819 | 0.206 |
| **7Networks_RH_Vis_9** | 4.023 | 0.000 | 4.050 | 0.000 | 2.271 | 0.012 | 0.844 | 0.199 | -0.447 | 0.328 | 0.727 | 0.234 |
| **7Networks_RH_Vis_12** | 2.908 | 0.002 | 2.994 | 0.001 | 1.398 | 0.081 | 1.320 | 0.094 | -2.069 | 0.019 | 1.882 | 0.030 |
| **7Networks_RH_Vis_14** | 3.667 | 0.000 | 3.005 | 0.001 | 3.075 | 0.001 | 2.659 | 0.004 | -2.209 | 0.014 | 2.729 | 0.003 |
| **7Networks_RH_SomMot_10** | 2.661 | 0.004 | 1.762 | 0.039 | 2.800 | 0.003 | -1.461 | 0.072 | 0.363 | 0.358 | -1.036 | 0.150 |
| **7Networks_RH_SomMot_12** | 2.660 | 0.004 | 1.981 | 0.024 | 2.414 | 0.008 | -0.714 | 0.238 | 0.330 | 0.371 | -0.589 | 0.278 |
| **7Networks_RH_SomMot_14** | 2.805 | 0.003 | 2.246 | 0.012 | 2.422 | 0.008 | -1.154 | 0.124 | 0.870 | 0.192 | -1.135 | 0.128 |
| **7Networks_RH_DorsAttn_PrCv_1** | 3.202 | 0.001 | 1.774 | 0.038 | 3.645 | 0.000 | 1.665 | 0.048 | 0.371 | 0.355 | 0.754 | 0.226 |
| **7Networks_RH_SalVentAttn_PrC_1** | 2.854 | 0.002 | 1.320 | 0.094 | 3.571 | 0.000 | 0.878 | 0.190 | -0.090 | 0.464 | 0.553 | 0.290 |
| **7Networks_RH_SalVentAttn_FrOperIns_2** | 3.479 | 0.000 | 2.619 | 0.004 | 3.520 | 0.000 | 1.004 | 0.158 | -0.820 | 0.206 | 1.022 | 0.153 |
| **7Networks_RH_Limbic_TempPole_1** | 4.514 | 0.000 | 3.827 | 0.000 | 3.257 | 0.001 | 0.682 | 0.248 | -1.843 | 0.033 | 1.392 | 0.082 |
| **7Networks_RH_Limbic_TempPole_2** | 3.540 | 0.000 | 2.981 | 0.001 | 2.648 | 0.004 | 1.669 | 0.048 | -0.899 | 0.184 | 1.447 | 0.074 |

| | | | | | | | | | | | | |
|---|---|---|---|---|---|---|---|---|---|---|---|---|
| **7Networks_RH_Limbic_TempPole_3** | 3.224 | 0.001 | 1.500 | 0.067 | 3.850 | 0.000 | 0.388 | 0.349 | -0.457 | 0.324 | 0.471 | 0.319 |
| **7Networks_RH_Cont_Temp_1** | 2.624 | 0.004 | 1.332 | 0.092 | 3.217 | 0.001 | 1.094 | 0.137 | -0.350 | 0.363 | 0.818 | 0.207 |
| **7Networks_RH_Cont_PFCl_1** | 3.043 | 0.001 | 2.182 | 0.015 | 2.935 | 0.002 | 0.930 | 0.176 | -0.412 | 0.340 | 0.758 | 0.224 |
| **7Networks_RH_Cont_Cing_2** | 2.630 | 0.004 | 2.644 | 0.004 | 1.634 | 0.051 | 0.947 | 0.172 | -0.114 | 0.455 | 0.606 | 0.272 |
| **7Networks_RH_Default_Temp_1** | 4.261 | 0.000 | 3.196 | 0.001 | 4.208 | 0.000 | 2.158 | 0.016 | -1.910 | 0.028 | 2.278 | 0.011 |
| **7Networks_RH_Default_Temp_2** | 3.212 | 0.001 | 1.476 | 0.070 | 4.229 | 0.000 | 2.523 | 0.006 | -0.784 | 0.217 | 1.874 | 0.031 |
| **7Networks_RH_Default_PFCdPFCm_7** | 4.332 | 0.000 | 3.432 | 0.000 | 3.500 | 0.000 | 0.540 | 0.295 | -0.313 | 0.377 | 0.480 | 0.316 |

***Supplementary Table 8.*** **Fluid cognition and local surface area.**

| | **Total Cognition** | **p** | **Fluid** | **p** | **Crystallized** | **p** | **Positive Affect** | **p** | **Negative Affect** | **p** | **Mean Affect** | **p** |
|---|---|---|---|---|---|---|---|---|---|---|---|---|
| **7Networks_LH_Vis_7** | 2.743 | 0.003 | 3.091 | 0.001 | 1.092 | 0.138 | 0.102 | 0.459 | -1.493 | 0.068 | 0.869 | 0.192 |
| **7Networks_LH_Vis_10** | 2.665 | 0.004 | 2.942 | 0.002 | 1.187 | 0.118 | 0.848 | 0.198 | -1.311 | 0.095 | 1.200 | 0.115 |
| **7Networks_LH_Vis_13** | 4.304 | 0.000 | 4.344 | 0.000 | 2.332 | 0.010 | 1.986 | 0.024 | -1.734 | 0.042 | 2.083 | 0.019 |
| **7Networks_LH_Vis_14** | 2.863 | 0.002 | 2.994 | 0.001 | 1.520 | 0.064 | 1.976 | 0.024 | -0.566 | 0.286 | 1.441 | 0.075 |
| **7Networks_LH_SomMot_12** | 3.113 | 0.001 | 2.916 | 0.002 | 2.259 | 0.012 | -0.318 | 0.375 | -0.357 | 0.360 | 0.011 | 0.495 |
| **7Networks_LH_Limbic_OFC_1** | 3.075 | 0.001 | 3.644 | 0.000 | 1.421 | 0.078 | 1.515 | 0.065 | -1.077 | 0.141 | 1.455 | 0.073 |
| **7Networks_LH_Limbic_TempPole_1** | 4.959 | 0.000 | 3.330 | 0.000 | 4.951 | 0.000 | 1.306 | 0.096 | -1.627 | 0.052 | 1.634 | 0.051 |
| **7Networks_RH_Vis_8** | 2.536 | 0.006 | 2.908 | 0.002 | 1.240 | 0.108 | 0.427 | 0.335 | -0.946 | 0.172 | 0.759 | 0.224 |
| **7Networks_RH_Vis_9** | 4.023 | 0.000 | 4.050 | 0.000 | 2.271 | 0.012 | 0.844 | 0.199 | -0.447 | 0.328 | 0.727 | 0.234 |
| **7Networks_RH_Vis_12** | 2.908 | 0.002 | 2.994 | 0.001 | 1.398 | 0.081 | 1.320 | 0.094 | -2.069 | 0.019 | 1.882 | 0.030 |
| **7Networks_RH_Vis_14** | 3.667 | 0.000 | 3.005 | 0.001 | 3.075 | 0.001 | 2.659 | 0.004 | -2.209 | 0.014 | 2.729 | 0.003 |
| **7Networks_RH_Limbic_TempPole_1** | 4.514 | 0.000 | 3.827 | 0.000 | 3.257 | 0.001 | 0.682 | 0.248 | -1.843 | 0.033 | 1.392 | 0.082 |
| **7Networks_RH_Limbic_TempPole_2** | 3.540 | 0.000 | 2.981 | 0.001 | 2.648 | 0.004 | 1.669 | 0.048 | -0.899 | 0.184 | 1.447 | 0.074 |
| **7Networks_RH_Default_Temp_1** | 4.261 | 0.000 | 3.196 | 0.001 | 4.208 | 0.000 | 2.158 | 0.016 | -1.910 | 0.028 | 2.278 | 0.011 |

*Supplementary Table 9.* **Crystallized cognition and local surface area.**

| | **Total Cognition** | **p** | **Fluid** | **p** | **Crystallized** | **p** | **Positive Affect** | **p** | **Negative Affect** | **p** | **Mean Affect** | **p** |
|---|---|---|---|---|---|---|---|---|---|---|---|---|
| **7Networks_LH_Vis_8** | 1.981 | 0.024 | 0.529 | 0.299 | 3.074 | 0.001 | -0.139 | 0.445 | -0.309 | 0.379 | 0.088 | 0.465 |
| **7Networks_LH_Vis_11** | 2.755 | 0.003 | 1.996 | 0.023 | 2.568 | 0.005 | 0.610 | 0.271 | -1.889 | 0.030 | 1.376 | 0.085 |
| **7Networks_LH_SomMot_4** | 1.403 | 0.080 | 0.066 | 0.474 | 2.557 | 0.005 | -0.123 | 0.451 | 0.858 | 0.195 | -0.537 | 0.296 |
| **7Networks_LH_SomMot_6** | 1.713 | 0.044 | 0.346 | 0.365 | 2.780 | 0.003 | -0.157 | 0.438 | -0.247 | 0.403 | 0.044 | 0.483 |
| **7Networks_LH_SomMot_8** | 3.059 | 0.001 | 2.224 | 0.013 | 2.947 | 0.002 | 1.751 | 0.040 | -1.605 | 0.054 | 1.878 | 0.030 |
| **7Networks_LH_SomMot_10** | 3.065 | 0.001 | 2.251 | 0.012 | 3.006 | 0.001 | -0.806 | 0.210 | 0.462 | 0.322 | -0.714 | 0.238 |
| **7Networks_LH_DorsAttn_Post_1** | 2.013 | 0.022 | 1.075 | 0.141 | 2.611 | 0.005 | -0.114 | 0.455 | 1.583 | 0.057 | -0.924 | 0.178 |
| **7Networks_LH_DorsAttn_Post_2** | 1.108 | 0.134 | -0.233 | 0.408 | 2.563 | 0.005 | -0.428 | 0.334 | 0.937 | 0.174 | -0.755 | 0.225 |
| **7Networks_LH_DorsAttn_Post_5** | 1.237 | 0.108 | -0.639 | 0.261 | 3.119 | 0.001 | 0.595 | 0.276 | 0.077 | 0.469 | 0.300 | 0.382 |
| **7Networks_LH_DorsAttn_PrCv_1** | 2.146 | 0.016 | 0.863 | 0.194 | 2.881 | 0.002 | 1.493 | 0.068 | -0.378 | 0.353 | 1.062 | 0.144 |
| **7Networks_LH_SalVentAttn_ParOper_1** | 2.061 | 0.020 | 0.863 | 0.194 | 2.523 | 0.006 | 1.212 | 0.113 | -0.071 | 0.472 | 0.734 | 0.232 |
| **7Networks_LH_SalVentAttn_FrOperIns_1** | 2.284 | 0.011 | 1.313 | 0.095 | 2.767 | 0.003 | 0.878 | 0.190 | -0.374 | 0.354 | 0.707 | 0.240 |
| **7Networks_LH_SalVentAttn_FrOperIns_3** | 1.653 | 0.049 | 0.485 | 0.314 | 2.698 | 0.004 | -0.146 | 0.442 | 0.173 | 0.431 | -0.178 | 0.429 |
| **7Networks_LH_SalVentAttn_FrOperIns_4** | 2.450 | 0.007 | 0.993 | 0.160 | 3.094 | 0.001 | 1.446 | 0.074 | -0.138 | 0.445 | 0.905 | 0.183 |
| **7Networks_LH_SalVentAttn_PFCl_1** | 2.444 | 0.007 | 0.546 | 0.293 | 3.838 | 0.000 | -1.347 | 0.089 | 1.071 | 0.142 | -1.355 | 0.088 |
| **7Networks_LH_Limbic_TempPole_1** | 4.959 | 0.000 | 3.330 | 0.000 | 4.951 | 0.000 | 1.306 | 0.096 | -1.627 | 0.052 | 1.634 | 0.051 |
| **7Networks_LH_Limbic_TempPole_2** | 3.216 | 0.001 | 2.079 | 0.019 | 3.297 | 0.001 | 1.862 | 0.031 | -1.545 | 0.061 | 1.909 | 0.028 |
| **7Networks_LH_Limbic_TempPole_3** | 4.238 | 0.000 | 2.830 | 0.002 | 4.458 | 0.000 | 1.039 | 0.149 | -0.603 | 0.273 | 0.924 | 0.178 |
| **7Networks_LH_Limbic_TempPole_4** | 3.346 | 0.000 | 2.135 | 0.017 | 3.898 | 0.000 | 0.620 | 0.268 | -0.210 | 0.417 | 0.470 | 0.319 |
| **7Networks_LH_Cont_PFCl_2** | 3.517 | 0.000 | 1.844 | 0.033 | 4.184 | 0.000 | 0.917 | 0.180 | -0.823 | 0.205 | 0.974 | 0.165 |
| **7Networks_LH_Cont_PFCl_5** | 3.376 | 0.000 | 1.633 | 0.051 | 4.074 | 0.000 | 0.224 | 0.411 | 1.590 | 0.056 | -0.734 | 0.232 |
| **7Networks_LH_Default_Temp_1** | 3.847 | 0.000 | 2.058 | 0.020 | 4.426 | 0.000 | 1.469 | 0.071 | -1.016 | 0.155 | 1.396 | 0.082 |
| **7Networks_LH_Default_Temp_2** | 4.388 | 0.000 | 2.659 | 0.004 | 4.597 | 0.000 | 1.333 | 0.091 | -1.205 | 0.114 | 1.420 | 0.078 |
| **7Networks_LH_Default_PFC_6** | 3.367 | 0.000 | 2.363 | 0.009 | 3.450 | 0.000 | 0.006 | 0.498 | -0.641 | 0.261 | 0.351 | 0.363 |
| **7Networks_LH_Default_PFC_8** | 2.630 | 0.004 | 1.816 | 0.035 | 2.689 | 0.004 | 0.018 | 0.493 | -0.626 | 0.266 | 0.350 | 0.363 |
| **7Networks_LH_Default_PFC_11** | 2.827 | 0.002 | 1.209 | 0.113 | 3.639 | 0.000 | 0.608 | 0.272 | 0.802 | 0.211 | -0.087 | 0.466 |

| | | | | | | | | | | | | |
|---|---|---|---|---|---|---|---|---|---|---|---|---|
| **7Networks_RH_Vis_1** | 2.359 | 0.009 | 1.135 | 0.128 | 2.764 | 0.003 | 2.083 | 0.019 | -0.965 | 0.167 | 1.720 | 0.043 |
| **7Networks_RH_Vis_4** | 3.155 | 0.001 | 2.349 | 0.009 | 2.774 | 0.003 | 0.991 | 0.161 | -0.460 | 0.323 | 0.819 | 0.206 |
| **7Networks_RH_Vis_5** | 2.397 | 0.008 | 1.285 | 0.100 | 3.050 | 0.001 | 0.400 | 0.345 | 0.101 | 0.460 | 0.175 | 0.431 |
| **7Networks_RH_Vis_14** | 3.667 | 0.000 | 3.005 | 0.001 | 3.075 | 0.001 | 2.659 | 0.004 | -2.209 | 0.014 | 2.729 | 0.003 |
| **7Networks_RH_SomMot_10** | 2.661 | 0.004 | 1.762 | 0.039 | 2.800 | 0.003 | -1.461 | 0.072 | 0.363 | 0.358 | -1.036 | 0.150 |
| **7Networks_RH_SomMot_13** | 2.376 | 0.009 | 0.794 | 0.214 | 3.175 | 0.001 | 1.257 | 0.105 | -1.470 | 0.071 | 1.520 | 0.064 |
| **7Networks_RH_SomMot_15** | 1.866 | 0.031 | 0.463 | 0.322 | 2.661 | 0.004 | 1.587 | 0.056 | -1.151 | 0.125 | 1.537 | 0.062 |
| **7Networks_RH_DorsAttn_Post_1** | 1.229 | 0.110 | -0.182 | 0.428 | 2.556 | 0.005 | 1.037 | 0.150 | 0.093 | 0.463 | 0.545 | 0.293 |
| **7Networks_RH_DorsAttn_PrCv_1** | 3.202 | 0.001 | 1.774 | 0.038 | 3.645 | 0.000 | 1.665 | 0.048 | 0.371 | 0.355 | 0.754 | 0.226 |
| **7Networks_RH_SalVentAttn_PrC_1** | 2.854 | 0.002 | 1.320 | 0.094 | 3.571 | 0.000 | 0.878 | 0.190 | -0.090 | 0.464 | 0.553 | 0.290 |
| **7Networks_RH_SalVentAttn_FrOperIns_1** | 2.251 | 0.012 | 1.111 | 0.134 | 3.197 | 0.001 | -0.409 | 0.341 | 0.754 | 0.226 | -0.644 | 0.260 |
| **7Networks_RH_SalVentAttn_FrOperIns_2** | 3.479 | 0.000 | 2.619 | 0.004 | 3.520 | 0.000 | 1.004 | 0.158 | -0.820 | 0.206 | 1.022 | 0.153 |
| **7Networks_RH_Limbic_OFC_3** | 2.387 | 0.009 | 1.144 | 0.126 | 2.897 | 0.002 | 1.893 | 0.029 | -1.225 | 0.110 | 1.752 | 0.040 |
| **7Networks_RH_Limbic_TempPole_1** | 4.514 | 0.000 | 3.827 | 0.000 | 3.257 | 0.001 | 0.682 | 0.248 | -1.843 | 0.033 | 1.392 | 0.082 |
| **7Networks_RH_Limbic_TempPole_2** | 3.540 | 0.000 | 2.981 | 0.001 | 2.648 | 0.004 | 1.669 | 0.048 | -0.899 | 0.184 | 1.447 | 0.074 |
| **7Networks_RH_Limbic_TempPole_3** | 3.224 | 0.001 | 1.500 | 0.067 | 3.850 | 0.000 | 0.388 | 0.349 | -0.457 | 0.324 | 0.471 | 0.319 |
| **7Networks_RH_Cont_Par_1** | 1.767 | 0.039 | 0.217 | 0.414 | 3.226 | 0.001 | 0.840 | 0.201 | -0.791 | 0.215 | 0.912 | 0.181 |
| **7Networks_RH_Cont_Par_2** | 1.089 | 0.138 | -0.739 | 0.230 | 3.079 | 0.001 | -0.744 | 0.229 | -0.422 | 0.337 | -0.198 | 0.422 |
| **7Networks_RH_Cont_Temp_1** | 2.624 | 0.004 | 1.332 | 0.092 | 3.217 | 0.001 | 1.094 | 0.137 | -0.350 | 0.363 | 0.818 | 0.207 |
| **7Networks_RH_Cont_PFCl_1** | 3.043 | 0.001 | 2.182 | 0.015 | 2.935 | 0.002 | 0.930 | 0.176 | -0.412 | 0.340 | 0.758 | 0.224 |
| **7Networks_RH_Cont_PFCl_2** | 1.216 | 0.112 | -0.262 | 0.397 | 2.774 | 0.003 | 1.293 | 0.098 | -0.992 | 0.161 | 1.281 | 0.100 |
| **7Networks_RH_Cont_PFCl_4** | 2.019 | 0.022 | 1.224 | 0.111 | 2.563 | 0.005 | -1.194 | 0.116 | 1.708 | 0.044 | -1.614 | 0.053 |
| **7Networks_RH_Cont_PFCl_5** | 0.813 | 0.208 | -0.760 | 0.224 | 2.551 | 0.005 | 0.653 | 0.257 | -0.185 | 0.427 | 0.476 | 0.317 |
| **7Networks_RH_Cont_pCun_1** | 2.068 | 0.019 | 1.283 | 0.100 | 2.520 | 0.006 | -0.007 | 0.497 | 0.561 | 0.288 | -0.309 | 0.379 |

***Supplementary Table 10.*** **Total cognition and subcortical volumes.**

| | **Total Cognition** | **p** | **Fluid** | **p** | **Crystallized** | **p** | **Positive Affect** | **p** | **Negative Affect** | **p** | **Mean Affect** | **p** |
|---|---|---|---|---|---|---|---|---|---|---|---|---|

| | | | | | | | | | | | | |
|---|---|---|---|---|---|---|---|---|---|---|---|---|
| **hipp_l** | 3.128 | 0.001 | 1.075 | 0.141 | 4.611 | 0.000 | -0.942 | 0.173 | 2.060 | 0.020 | -1.660 | 0.049 |

***Supplementary Table 11.*** **Fluid cognition and subcortical volumes.**

| | Total Cognition | p | Fluid | p | Crystallized | p | Positive Affect | p | Negative Affect | p | Mean Affect | p |
|---|---|---|---|---|---|---|---|---|---|---|---|---|
| **pall_l** | -2.669 | 0.004 | -3.147 | 0.001 | -0.922 | 0.178 | -0.465 | 0.321 | 1.432 | 0.076 | -1.045 | 0.148 |

***Supplementary Table 12.*** **Crystallized cognition and subcortical volumes.**

| | Total Cognition | p | Fluid | p | Crystallized | p | Positive Affect | p | Negative Affect | p | Mean Affect | p |
|---|---|---|---|---|---|---|---|---|---|---|---|---|
| **amy_r** | 1.329 | 0.092 | -0.045 | 0.482 | 2.738 | 0.003 | -0.023 | 0.491 | 1.402 | 0.081 | -0.774 | 0.219 |
| **hipp_l** | 3.128 | 0.001 | 1.075 | 0.141 | 4.611 | 0.000 | -0.942 | 0.173 | 2.060 | 0.020 | -1.660 | 0.049 |
| **hipp_r** | 2.687 | 0.004 | 0.870 | 0.192 | 4.063 | 0.000 | -0.056 | 0.477 | 1.587 | 0.056 | -0.894 | 0.186 |

***Supplementary Table 13.*** **Mean affect and subcortical volumes.**

| | Total Cognition | p | Fluid | p | Crystallized | p | Positive Affect | p | Negative Affect | p | Mean Affect | p |
|---|---|---|---|---|---|---|---|---|---|---|---|---|
| **caud_l** | 0.082 | 0.467 | -1.050 | 0.147 | 1.595 | 0.056 | -4.039 | 0.000 | 3.497 | 0.000 | -4.226 | 0.000 |
| **caud_r** | 0.082 | 0.467 | -0.991 | 0.161 | 1.499 | 0.067 | -3.347 | 0.000 | 3.175 | 0.001 | -3.651 | 0.000 |

***Supplementary Table 14.*** **Positive affect and subcortical volumes.**

| | Total Cognition | p | Fluid | p | Crystallized | p | Positive Affect | p | Negative Affect | p | Mean Affect | p |
|---|---|---|---|---|---|---|---|---|---|---|---|---|
| **caud_l** | 0.082 | 0.467 | -1.050 | 0.147 | 1.595 | 0.056 | -4.039 | 0.000 | 3.497 | 0.000 | -4.226 | 0.000 |
| **caud_r** | 0.082 | 0.467 | -0.991 | 0.161 | 1.499 | 0.067 | -3.347 | 0.000 | 3.175 | 0.001 | -3.651 | 0.000 |

***Supplementary Table 15.*** **Negative affect and subcortical volumes.**

| | **Total Cognition** | **p** | **Fluid** | **p** | **Crystallized** | **p** | **Positive Affect** | **p** | **Negative Affect** | **p** | **Mean Affect** | **p** |
|---|---|---|---|---|---|---|---|---|---|---|---|---|
| **caud_l** | 0.082 | 0.467 | -1.050 | 0.147 | 1.595 | 0.056 | -4.039 | 0.000 | 3.497 | 0.000 | -4.226 | 0.000 |
| **caud_r** | 0.082 | 0.467 | -0.991 | 0.161 | 1.499 | 0.067 | -3.347 | 0.000 | 3.175 | 0.001 | -3.651 | 0.000 |

***Supplementary Table 16.* Genetic correlation of cognition and affect with local cortical thickness.** Environmental correlation ($\rho_e$) and genetic correlation ($\rho_g$) per parcel associated with total cognition score and mean affect score, respectively (see Figure 3A).

| **Total Cognition** | $\rho_e$ | **p** | $\rho_g$ | **p** |
|---:|---:|---:|---:|---:|
| *7Networks_LH_Vis_1* | -0.027 | 0.697 | 0.211 | 0.003 |
| *7Networks_LH_Vis_4* | 0.000 | 0.998 | 0.182 | 0.004 |
| *7Networks_LH_Vis_9* | -0.057 | 0.412 | 0.189 | 0.003 |
| *7Networks_LH_Vis_10* | 0.029 | 0.688 | 0.134 | 0.020 |
| *7Networks_LH_SomMot_3* | 0.009 | 0.905 | 0.201 | 0.004 |
| *7Networks_LH_SomMot_10* | -0.078 | 0.265 | 0.327 | 0.000 |
| *7Networks_LH_DorsAttn_Post_7* | 0.144 | 0.052 | -0.235 | 0.001 |
| *7Networks_LH_DorsAttn_FEF_2* | 0.043 | 0.559 | -0.214 | 0.009 |
| *7Networks_LH_SalVentAttn_PFCl_1* | 0.113 | 0.096 | -0.274 | 0.000 |
| *7Networks_LH_Default_Temp_5* | -0.046 | 0.528 | 0.245 | 0.014 |
| *7Networks_LH_Default_PFC_4* | 0.018 | 0.806 | -0.177 | 0.007 |
| *7Networks_LH_Default_PFC_5* | 0.124 | 0.084 | -0.281 | 0.001 |
| *7Networks_LH_Default_PFC_7* | -0.018 | 0.802 | -0.212 | 0.001 |
| *7Networks_LH_Default_PFC_9* | 0.017 | 0.814 | -0.171 | 0.007 |
| *7Networks_LH_Default_PFC_10* | -0.044 | 0.530 | -0.235 | 0.002 |
| *7Networks_LH_Default_PFC_11* | 0.124 | 0.075 | -0.432 | 0.000 |
| *7Networks_LH_Default_PFC_13* | 0.042 | 0.567 | -0.212 | 0.003 |
| *7Networks_LH_Default_pCunPCC_1* | 0.014 | 0.852 | 0.187 | 0.008 |
| *7Networks_RH_Vis_4* | -0.119 | 0.104 | 0.176 | 0.004 |
| *7Networks_RH_Vis_9* | -0.075 | 0.310 | 0.158 | 0.009 |
| *7Networks_RH_Vis_10* | -0.173 | 0.018 | 0.256 | 0.000 |
| *7Networks_RH_Vis_13* | -0.091 | 0.209 | 0.215 | 0.001 |
| *7Networks_RH_SomMot_7* | -0.023 | 0.754 | 0.137 | 0.035 |
| *7Networks_RH_SomMot_8* | -0.053 | 0.464 | 0.256 | 0.003 |
| *7Networks_RH_SomMot_12* | -0.060 | 0.403 | 0.226 | 0.001 |
| *7Networks_RH_Cont_PFCv_1* | -0.106 | 0.127 | 0.313 | 0.001 |
| *7Networks_RH_Cont_PFCl_5* | -0.011 | 0.875 | -0.179 | 0.015 |
| *7Networks_RH_Cont_PFCl_6* | -0.008 | 0.909 | -0.210 | 0.007 |
| *7Networks_RH_Cont_PFCl_7* | 0.140 | 0.049 | -0.287 | 0.001 |
| *7Networks_RH_Cont_PFCmp_2* | 0.179 | 0.016 | -0.283 | 0.000 |
| *7Networks_RH_Default_PFCdPFCm_4* | 0.051 | 0.482 | -0.247 | 0.000 |
| *7Networks_RH_Default_PFCdPFCm_5* | -0.078 | 0.283 | -0.217 | 0.001 |
| *7Networks_RH_Default_PFCdPFCm_6* | 0.150 | 0.036 | -0.299 | 0.000 |
| *7Networks_RH_Default_pCunPCC_1* | -0.043 | 0.563 | 0.329 | 0.000 |
| **Mean affect** | $\rho_e$ | **p** | $\rho_g$ | **p** |
| *7Networks_LH_Default_PFC_9* | 0.120 | 0.071 | -0.480 | 0.000 |

***Supplementary Table 17.*** **Genetic correlation between cognition and local surface area.** Environmental correlation ($\rho_e$) and genetic correlation ($\rho_g$) per parcel associated with total cognition score (see Figure 3B).

| **Total Cognition** | $\rho_e$ | **p** | $\rho_g$ | **p** |
|---|---|---|---|---|
| *7Networks_LH_Vis_7* | 0.025 | 0.742 | 0.118 | 0.042 |
| *7Networks_LH_Vis_10* | -0.063 | 0.411 | 0.101 | 0.039 |
| *7Networks_LH_Vis_13* | -0.128 | 0.085 | 0.239 | 0.000 |
| *7Networks_LH_Vis_14* | 0.022 | 0.770 | 0.150 | 0.034 |
| *7Networks_LH_SomMot_8* | 0.000 | 0.997 | 0.151 | 0.024 |
| *7Networks_LH_SomMot_10* | 0.039 | 0.564 | 0.172 | 0.024 |
| *7Networks_LH_SomMot_12* | 0.003 | 0.966 | 0.182 | 0.006 |
| *7Networks_LH_SalVentAttn_Med_2* | -0.074 | 0.303 | 0.206 | 0.004 |
| *7Networks_LH_Limbic_TempPole_1* | 0.008 | 0.911 | 0.213 | 0.001 |
| *7Networks_LH_Limbic_TempPole_3* | 0.050 | 0.474 | 0.174 | 0.015 |
| *7Networks_LH_Limbic_TempPole_4* | -0.030 | 0.682 | 0.167 | 0.006 |
| *7Networks_LH_Cont_PFCl_2* | -0.064 | 0.387 | 0.242 | 0.001 |
| *7Networks_LH_Cont_PFCl_5* | 0.055 | 0.421 | 0.205 | 0.037 |
| *7Networks_LH_Default_Temp_1* | -0.111 | 0.141 | 0.199 | 0.001 |
| *7Networks_LH_Default_Temp_2* | -0.044 | 0.536 | 0.293 | 0.000 |
| *7Networks_LH_Default_PFC_6* | -0.017 | 0.823 | 0.154 | 0.011 |
| *7Networks_LH_Default_PFC_8* | -0.068 | 0.350 | 0.179 | 0.012 |
| *7Networks_RH_Vis_4* | 0.009 | 0.904 | 0.135 | 0.037 |
| *7Networks_RH_Vis_9* | -0.068 | 0.390 | 0.176 | 0.001 |
| *7Networks_RH_Vis_12* | -0.004 | 0.954 | 0.147 | 0.017 |
| *7Networks_RH_Vis_14* | -0.070 | 0.323 | 0.257 | 0.001 |
| *7Networks_RH_Limbic_TempPole_1* | -0.204 | 0.004 | 0.322 | 0.000 |
| *7Networks_RH_Limbic_TempPole_2* | -0.119 | 0.094 | 0.231 | 0.001 |
| *7Networks_RH_Limbic_TempPole_3* | -0.076 | 0.290 | 0.202 | 0.004 |
| *7Networks_RH_Cont_PFCl_1* | -0.017 | 0.805 | 0.163 | 0.009 |
| *7Networks_RH_Cont_Cing_2* | -0.103 | 0.153 | 0.210 | 0.006 |
| *7Networks_RH_Default_Temp_1* | 0.073 | 0.291 | 0.173 | 0.008 |
| *7Networks_RH_Default_Temp_2* | -0.042 | 0.549 | 0.193 | 0.009 |
| *7Networks_RH_Default_PFCdPFCm_7* | 0.030 | 0.666 | 0.228 | 0.008 |

***Supplementary Table 18.*** **Genetic correlation of cognitive and affective sub-scores and local cortical thickness.**

| **Fluid Cognition** | $\rho_e$ | **p** | $\rho_g$ | **p** |
|---|---|---|---|---|
| *7Networks_LH_Vis_9* | -0.058 | 0.388 | 0.240 | 0.002 |
| *7Networks_LH_Vis_10* | 0.048 | 0.498 | 0.146 | 0.039 |
| *7Networks_LH_SomMot_10* | -0.069 | 0.300 | 0.319 | 0.000 |
| *7Networks_LH_SalVentAttn_PFCl_1* | 0.131 | 0.043 | -0.315 | 0.000 |
| *7Networks_LH_Default_PFC_7* | 0.043 | 0.525 | -0.258 | 0.002 |
| *7Networks_LH_Default_PFC_9* | 0.066 | 0.343 | -0.240 | 0.002 |
| *7Networks_LH_Default_PFC_10* | 0.056 | 0.401 | -0.355 | 0.000 |

| | | | | |
|---|---|---|---|---|
| *7Networks_LH_Default_PFC_11* | 0.145 | 0.026 | -0.479 | 0.000 |
| *7Networks_LH_Default_PFC_13* | 0.049 | 0.471 | -0.271 | 0.002 |
| *7Networks_RH_Vis_9* | -0.133 | 0.060 | 0.270 | 0.000 |
| *7Networks_RH_Vis_10* | -0.177 | 0.011 | 0.333 | 0.000 |
| *7Networks_RH_Vis_13* | -0.127 | 0.066 | 0.287 | 0.000 |
| *7Networks_RH_SomMot_12* | -0.089 | 0.191 | 0.298 | 0.001 |
| *7Networks_RH_Cont_PFCmp_2* | 0.195 | 0.005 | -0.362 | 0.000 |
| *7Networks_RH_Default_PFCdPFCm_4* | 0.060 | 0.389 | -0.231 | 0.005 |
| *7Networks_RH_Default_PFCdPFCm_5* | -0.022 | 0.751 | -0.260 | 0.001 |
| *7Networks_RH_Default_PFCdPFCm_6* | 0.166 | 0.015 | -0.380 | 0.000 |
| *7Networks_RH_Default_pCunPCC_1* | -0.044 | 0.531 | 0.308 | 0.001 |
| **Crystallized Cognition** | $\rho_e$ | **p** | $\rho_g$ | **p** |
| *7Networks_LH_Vis_1* | 0.038 | 0.609 | 0.220 | 0.001 |
| *7Networks_LH_Vis_2* | 0.053 | 0.471 | 0.178 | 0.007 |
| *7Networks_LH_Vis_4* | -0.009 | 0.896 | 0.184 | 0.001 |
| *7Networks_LH_SomMot_3* | -0.001 | 0.987 | 0.205 | 0.001 |
| *7Networks_LH_SomMot_10* | -0.097 | 0.185 | 0.287 | 0.000 |
| *7Networks_LH_DorsAttn_Post_1* | -0.084 | 0.291 | 0.204 | 0.004 |
| *7Networks_LH_DorsAttn_Post_7* | 0.087 | 0.270 | -0.213 | 0.001 |
| *7Networks_LH_DorsAttn_Post_9* | 0.015 | 0.839 | -0.135 | 0.021 |
| *7Networks_LH_SalVentAttn_FrOperIns_2* | 0.050 | 0.509 | 0.131 | 0.045 |
| *7Networks_LH_SalVentAttn_FrOperIns_3* | -0.044 | 0.566 | 0.173 | 0.010 |
| *7Networks_LH_SalVentAttn_PFCl_1* | 0.096 | 0.177 | -0.219 | 0.000 |
| *7Networks_LH_Cont_PFCl_2* | 0.082 | 0.274 | -0.189 | 0.003 |
| *7Networks_LH_Cont_PFCl_3* | -0.011 | 0.885 | -0.194 | 0.013 |
| *7Networks_LH_Default_PFC_4* | 0.018 | 0.816 | -0.137 | 0.020 |
| *7Networks_LH_Default_PFC_7* | -0.048 | 0.523 | -0.160 | 0.007 |
| *7Networks_LH_Default_PFC_11* | 0.069 | 0.353 | -0.312 | 0.000 |
| *7Networks_LH_Default_pCunPCC_1* | -0.039 | 0.607 | 0.177 | 0.006 |
| *7Networks_RH_Vis_3* | -0.084 | 0.273 | 0.204 | 0.001 |
| *7Networks_RH_Vis_4* | -0.067 | 0.380 | 0.220 | 0.000 |
| *7Networks_RH_Vis_8* | -0.007 | 0.926 | 0.162 | 0.006 |
| *7Networks_RH_SomMot_1* | 0.053 | 0.492 | 0.144 | 0.020 |
| *7Networks_RH_SomMot_3* | -0.049 | 0.535 | 0.188 | 0.007 |
| *7Networks_RH_SomMot_8* | 0.011 | 0.884 | 0.214 | 0.006 |
| *7Networks_RH_SomMot_10* | -0.027 | 0.724 | 0.205 | 0.029 |
| *7Networks_RH_SalVentAttn_TempOccPar_1* | -0.018 | 0.814 | 0.188 | 0.016 |
| *7Networks_RH_SalVentAttn_FrOperIns_4* | 0.008 | 0.917 | 0.193 | 0.024 |
| *7Networks_RH_Limbic_TempPole_1* | -0.011 | 0.883 | 0.210 | 0.011 |
| *7Networks_RH_Cont_PFCv_1* | -0.085 | 0.250 | 0.249 | 0.003 |
| *7Networks_RH_Cont_PFCl_3* | -0.035 | 0.646 | -0.149 | 0.025 |
| *7Networks_RH_Cont_PFCl_5* | 0.030 | 0.681 | -0.186 | 0.005 |
| *7Networks_RH_Cont_PFCl_6* | -0.112 | 0.117 | -0.167 | 0.016 |
| *7Networks_RH_Default_Par_1* | 0.040 | 0.610 | -0.251 | 0.009 |

| | | | | |
|---|---|---|---|---|
| *7Networks_RH_Default_Temp_5* | -0.035 | 0.643 | 0.299 | 0.001 |
| *7Networks_RH_Default_PFCdPFCm_4* | 0.017 | 0.830 | -0.208 | 0.001 |
| *7Networks_RH_Default_PFCdPFCm_5* | -0.094 | 0.213 | -0.135 | 0.020 |
| *7Networks_RH_Default_pCunPCC_1* | -0.030 | 0.706 | 0.280 | 0.000 |
| **Positive affect** | $\rho_e$ | **p** | $\rho_g$ | **p** |
| *7Networks_LH_Default_PFC_9* | 0.129 | 0.044 | -0.540 | 0.000 |
| **Negative affect** | $\rho_e$ | **p** | $\rho_g$ | **p** |
| *7Networks_LH_Vis_14* | -0.211 | 0.002 | 0.011 | 0.925 |

***Supplementary Table 19.*** **Genetic correlation of cognitive sub-scores and local surface area.**

| **Fluid Cognition** | $\rho_e$ | **p** | $\rho_g$ | **p** |
|---|---|---|---|---|
| *7Networks_LH_Vis_7* | 0.006 | 0.937 | 0.158 | 0.028 |
| *7Networks_LH_Vis_10* | -0.085 | 0.268 | 0.158 | 0.009 |
| *7Networks_LH_Vis_13* | -0.109 | 0.132 | 0.290 | 0.000 |
| *7Networks_LH_SomMot_12* | 0.032 | 0.631 | 0.165 | 0.042 |
| *7Networks_LH_Limbic_TempPole_1* | -0.018 | 0.796 | 0.182 | 0.023 |
| *7Networks_RH_Vis_9* | -0.082 | 0.296 | 0.222 | 0.000 |
| *7Networks_RH_Vis_12* | -0.005 | 0.944 | 0.182 | 0.018 |
| *7Networks_RH_Vis_14* | -0.083 | 0.215 | 0.284 | 0.002 |
| *7Networks_RH_Limbic_TempPole_1* | -0.179 | 0.008 | 0.361 | 0.000 |
| *7Networks_RH_Limbic_TempPole_2* | -0.156 | 0.020 | 0.311 | 0.000 |
| *7Networks_RH_Default_Temp_1* | 0.001 | 0.985 | 0.179 | 0.028 |
| **Crystallized Cognition** | $\rho_e$ | **p** | $\rho_g$ | **p** |
| *7Networks_LH_SomMot_4* | -0.058 | 0.433 | 0.168 | 0.022 |
| *7Networks_LH_SomMot_6* | -0.041 | 0.577 | 0.147 | 0.011 |
| *7Networks_LH_DorsAttn_Post_1* | -0.121 | 0.124 | 0.197 | 0.003 |
| *7Networks_LH_DorsAttn_Post_2* | -0.068 | 0.378 | 0.175 | 0.008 |
| *7Networks_LH_DorsAttn_Post_5* | -0.014 | 0.852 | 0.221 | 0.010 |
| *7Networks_LH_DorsAttn_PrCv_1* | 0.003 | 0.972 | 0.176 | 0.016 |
| *7Networks_LH_SalVentAttn_PFCl_1* | -0.172 | 0.035 | 0.324 | 0.000 |
| *7Networks_LH_Limbic_TempPole_1* | 0.024 | 0.757 | 0.203 | 0.000 |
| *7Networks_LH_Limbic_TempPole_2* | 0.019 | 0.797 | 0.136 | 0.026 |
| *7Networks_LH_Limbic_TempPole_3* | 0.060 | 0.409 | 0.180 | 0.005 |
| *7Networks_LH_Limbic_TempPole_4* | -0.058 | 0.451 | 0.190 | 0.000 |
| *7Networks_LH_Cont_PFCl_2* | -0.069 | 0.379 | 0.255 | 0.000 |
| *7Networks_LH_Cont_PFCl_5* | 0.089 | 0.220 | 0.223 | 0.010 |
| *7Networks_LH_Default_Temp_1* | 0.016 | 0.837 | 0.155 | 0.002 |
| *7Networks_LH_Default_Temp_2* | -0.052 | 0.493 | 0.282 | 0.000 |
| *7Networks_LH_Default_PFC_6* | 0.066 | 0.394 | 0.108 | 0.043 |
| *7Networks_RH_Vis_1* | -0.083 | 0.258 | 0.145 | 0.010 |
| *7Networks_RH_Vis_5* | -0.136 | 0.066 | 0.298 | 0.001 |

| | | | | |
|---|---|---|---|---|
| *7Networks_RH_Vis_14* | -0.002 | 0.977 | 0.161 | 0.015 |
| *7Networks_RH_SomMot_13* | -0.048 | 0.538 | 0.200 | 0.005 |
| *7Networks_RH_DorsAttn_Post_1* | -0.034 | 0.657 | 0.117 | 0.045 |
| *7Networks_RH_DorsAttn_PrCv_1* | 0.011 | 0.882 | 0.221 | 0.002 |
| *7Networks_RH_SalVentAttn_PrC_1* | 0.054 | 0.472 | 0.202 | 0.008 |
| *7Networks_RH_Limbic_OFC_3* | 0.000 | 0.996 | 0.137 | 0.032 |
| *7Networks_RH_Limbic_TempPole_1* | -0.118 | 0.106 | 0.183 | 0.001 |
| *7Networks_RH_Limbic_TempPole_3* | -0.069 | 0.358 | 0.204 | 0.001 |
| *7Networks_RH_Cont_Par_2* | 0.006 | 0.938 | 0.200 | 0.026 |
| *7Networks_RH_Cont_Temp_1* | -0.025 | 0.756 | 0.180 | 0.012 |
| *7Networks_RH_Cont_PFCl_1* | 0.042 | 0.568 | 0.117 | 0.035 |
| *7Networks_RH_Cont_PFCl_2* | -0.056 | 0.473 | 0.174 | 0.019 |
| *7Networks_RH_Cont_PFCmp_2* | -0.108 | 0.146 | 0.180 | 0.004 |
| *7Networks_RH_Default_Temp_1* | 0.135 | 0.059 | 0.153 | 0.008 |
| *7Networks_RH_Default_Temp_2* | 0.115 | 0.112 | 0.150 | 0.019 |
| *7Networks_RH_Default_Temp_4* | 0.023 | 0.749 | 0.167 | 0.013 |
| *7Networks_RH_Default_PFCdPFCm_6* | -0.092 | 0.213 | 0.195 | 0.003 |
| *7Networks_RH_Default_PFCdPFCm_7* | -0.134 | 0.072 | 0.269 | 0.000 |

***Supplementary Table 20.*** **Genetic correlation of cognitive and affective scores and subcortical volumes.**

| **Total Cognition** | $\rho_e$ | **p** | $\rho_g$ | **p** |
|---|---|---|---|---|
| *hipp_l* | 0.175 | 0.015 | 0.057 | 0.349 |
| **Fluid Cognition** | $\rho_e$ | **p** | $\rho_g$ | **p** |
| *pall_l* | 0.089 | 0.212 | -0.230 | 0.003 |
| **Crystallized Cognition** | $\rho_e$ | **p** | $\rho_g$ | **p** |
| *amy_r* | 0.042 | 0.621 | 0.075 | 0.138 |
| *hipp_l* | 0.124 | 0.093 | 0.159 | 0.004 |
| *hipp_r* | 0.214 | 0.008 | 0.098 | 0.026 |
| **Mean affect** | $\rho_e$ | **p** | $\rho_g$ | **p** |
| *caud_l* | 0.041 | 0.603 | -0.280 | 0.001 |
| *caud_r* | 0.084 | 0.297 | -0.283 | 0.001 |
| **Positive affect** | $\rho_e$ | **p** | $\rho_g$ | **p** |
| *caud_l* | 0.018 | 0.820 | -0.275 | 0.002 |
| *caud_r* | 0.083 | 0.284 | -0.287 | 0.001 |
| **Negative affect** | $\rho_e$ | **p** | $\rho_g$ | **p** |
| *caud_l* | -0.055 | 0.499 | 0.224 | 0.003 |
| *caud_r* | -0.052 | 0.526 | 0.217 | 0.005 |

***Supplementary Table 21.* Heritability of local cortical thickness.**

| Parcel | $h^2$ | p |
|---|---|---|
| 7Networks_LH_Vis_1 | 0.411 | 0.000 |
| 7Networks_LH_Vis_2 | 0.387 | 0.000 |
| 7Networks_LH_Vis_3 | 0.168 | 0.002 |
| 7Networks_LH_Vis_4 | 0.476 | 0.000 |
| 7Networks_LH_Vis_5 | 0.512 | 0.000 |
| 7Networks_LH_Vis_6 | 0.437 | 0.000 |
| 7Networks_LH_Vis_7 | 0.509 | 0.000 |
| 7Networks_LH_Vis_8 | 0.237 | 0.000 |
| 7Networks_LH_Vis_9 | 0.472 | 0.000 |
| 7Networks_LH_Vis_10 | 0.600 | 0.000 |
| 7Networks_LH_Vis_11 | 0.400 | 0.000 |
| 7Networks_LH_Vis_12 | 0.464 | 0.000 |
| 7Networks_LH_Vis_13 | 0.557 | 0.000 |
| 7Networks_LH_Vis_14 | 0.454 | 0.000 |
| 7Networks_LH_SomMot_1 | 0.445 | 0.000 |
| 7Networks_LH_SomMot_2 | 0.260 | 0.000 |
| 7Networks_LH_SomMot_3 | 0.439 | 0.000 |
| 7Networks_LH_SomMot_4 | 0.416 | 0.000 |
| 7Networks_LH_SomMot_5 | 0.274 | 0.000 |
| 7Networks_LH_SomMot_6 | 0.511 | 0.000 |
| 7Networks_LH_SomMot_7 | 0.330 | 0.000 |
| 7Networks_LH_SomMot_8 | 0.269 | 0.000 |
| 7Networks_LH_SomMot_9 | 0.305 | 0.000 |
| 7Networks_LH_SomMot_10 | 0.460 | 0.000 |
| 7Networks_LH_SomMot_11 | 0.199 | 0.000 |
| 7Networks_LH_SomMot_12 | 0.423 | 0.000 |
| 7Networks_LH_SomMot_13 | 0.367 | 0.000 |
| 7Networks_LH_SomMot_14 | 0.332 | 0.000 |
| 7Networks_LH_SomMot_15 | 0.531 | 0.000 |
| 7Networks_LH_SomMot_16 | 0.348 | 0.000 |
| 7Networks_LH_DorsAttn_Post_1 | 0.364 | 0.000 |
| 7Networks_LH_DorsAttn_Post_2 | 0.240 | 0.000 |
| 7Networks_LH_DorsAttn_Post_3 | 0.349 | 0.000 |
| 7Networks_LH_DorsAttn_Post_4 | 0.298 | 0.000 |
| 7Networks_LH_DorsAttn_Post_5 | 0.142 | 0.004 |
| 7Networks_LH_DorsAttn_Post_6 | 0.225 | 0.000 |
| 7Networks_LH_DorsAttn_Post_7 | 0.460 | 0.000 |
| 7Networks_LH_DorsAttn_Post_8 | 0.278 | 0.000 |
| 7Networks_LH_DorsAttn_Post_9 | 0.486 | 0.000 |
| 7Networks_LH_DorsAttn_Post_10 | 0.479 | 0.000 |
| 7Networks_LH_DorsAttn_FEF_1 | 0.307 | 0.000 |
| 7Networks_LH_DorsAttn_FEF_2 | 0.337 | 0.000 |
| 7Networks_LH_DorsAttn_PrCv_1 | 0.272 | 0.000 |
| 7Networks_LH_SalVentAttn_ParOper_1 | 0.102 | 0.043 |
| 7Networks_LH_SalVentAttn_ParOper_2 | 0.311 | 0.000 |
| 7Networks_LH_SalVentAttn_ParOper_3 | 0.255 | 0.000 |
| 7Networks_LH_SalVentAttn_FrOperIns_1 | 0.309 | 0.000 |
| 7Networks_LH_SalVentAttn_FrOperIns_2 | 0.404 | 0.000 |
| 7Networks_LH_SalVentAttn_FrOperIns_3 | 0.392 | 0.000 |
| 7Networks_LH_SalVentAttn_FrOperIns_4 | 0.225 | 0.000 |
| 7Networks_LH_SalVentAttn_PFCl_1 | 0.434 | 0.000 |
| 7Networks_LH_SalVentAttn_Med_1 | 0.343 | 0.000 |
| 7Networks_LH_SalVentAttn_Med_2 | 0.351 | 0.000 |
| 7Networks_LH_SalVentAttn_Med_3 | 0.288 | 0.000 |
| 7Networks_LH_Limbic_OFC_1 | 0.306 | 0.000 |
| 7Networks_LH_Limbic_OFC_2 | 0.423 | 0.000 |
| 7Networks_LH_Limbic_TempPole_1 | 0.479 | 0.000 |
| 7Networks_LH_Limbic_TempPole_2 | 0.356 | 0.000 |
| 7Networks_LH_Limbic_TempPole_3 | 0.241 | 0.000 |
| 7Networks_LH_Limbic_TempPole_4 | 0.430 | 0.000 |
| 7Networks_LH_Cont_Par_1 | 0.257 | 0.000 |
| 7Networks_LH_Cont_Par_2 | 0.288 | 0.000 |
| 7Networks_LH_Cont_Par_3 | 0.113 | 0.022 |
| 7Networks_LH_Cont_Temp_1 | 0.203 | 0.000 |
| 7Networks_LH_Cont_OFC_1 | 0.348 | 0.000 |
| 7Networks_LH_Cont_PFCl_1 | 0.280 | 0.000 |
| 7Networks_LH_Cont_PFCl_2 | 0.404 | 0.000 |
| 7Networks_LH_Cont_PFCl_3 | 0.274 | 0.000 |
| 7Networks_LH_Cont_PFCl_4 | 0.373 | 0.000 |
| 7Networks_LH_Cont_PFCl_5 | 0.234 | 0.000 |
| 7Networks_LH_Cont_pCun_1 | 0.387 | 0.000 |
| 7Networks_LH_Cont_Cing_1 | 0.558 | 0.000 |
| 7Networks_LH_Cont_Cing_2 | 0.427 | 0.000 |
| 7Networks_LH_Default_Temp_1 | 0.250 | 0.000 |
| 7Networks_LH_Default_Temp_2 | 0.290 | 0.000 |
| 7Networks_LH_Default_Temp_3 | 0.350 | 0.000 |
| 7Networks_LH_Default_Temp_4 | 0.226 | 0.000 |
| 7Networks_LH_Default_Temp_5 | 0.238 | 0.000 |
| 7Networks_LH_Default_Par_1 | 0.330 | 0.000 |
| 7Networks_LH_Default_Par_2 | 0.328 | 0.000 |
| 7Networks_LH_Default_Par_3 | 0.240 | 0.000 |
| 7Networks_LH_Default_Par_4 | 0.255 | 0.000 |
| 7Networks_LH_Default_PFC_1 | 0.355 | 0.000 |
| 7Networks_LH_Default_PFC_2 | 0.323 | 0.000 |

| | | |
|---|---|---|
| 7Networks_LH_Default_PFC_3 | 0.168 | 0.001 |
| 7Networks_LH_Default_PFC_4 | 0.488 | 0.000 |
| 7Networks_LH_Default_PFC_5 | 0.322 | 0.000 |
| 7Networks_LH_Default_PFC_6 | 0.489 | 0.000 |
| 7Networks_LH_Default_PFC_7 | 0.483 | 0.000 |
| 7Networks_LH_Default_PFC_8 | 0.340 | 0.000 |
| 7Networks_LH_Default_PFC_9 | 0.517 | 0.000 |
| 7Networks_LH_Default_PFC_10 | 0.359 | 0.000 |
| 7Networks_LH_Default_PFC_11 | 0.443 | 0.000 |
| 7Networks_LH_Default_PFC_12 | 0.471 | 0.000 |
| 7Networks_LH_Default_PFC_13 | 0.429 | 0.000 |
| 7Networks_LH_Default_pCunPCC_1 | 0.434 | 0.000 |
| 7Networks_LH_Default_pCunPCC_2 | 0.240 | 0.000 |
| 7Networks_LH_Default_pCunPCC_3 | 0.458 | 0.000 |
| 7Networks_LH_Default_pCunPCC_4 | 0.362 | 0.000 |
| 7Networks_LH_Default_PHC_1 | 0.484 | 0.000 |
| 7Networks_RH_Vis_1 | 0.379 | 0.000 |
| 7Networks_RH_Vis_2 | 0.437 | 0.000 |
| 7Networks_RH_Vis_3 | 0.436 | 0.000 |
| 7Networks_RH_Vis_4 | 0.580 | 0.000 |
| 7Networks_RH_Vis_5 | 0.336 | 0.000 |
| 7Networks_RH_Vis_6 | 0.544 | 0.000 |
| 7Networks_RH_Vis_7 | 0.523 | 0.000 |
| 7Networks_RH_Vis_8 | 0.505 | 0.000 |
| 7Networks_RH_Vis_9 | 0.585 | 0.000 |
| 7Networks_RH_Vis_10 | 0.514 | 0.000 |
| 7Networks_RH_Vis_11 | 0.277 | 0.000 |
| 7Networks_RH_Vis_12 | 0.544 | 0.000 |
| 7Networks_RH_Vis_13 | 0.520 | 0.000 |
| 7Networks_RH_Vis_14 | 0.345 | 0.000 |
| 7Networks_RH_Vis_15 | 0.342 | 0.000 |
| 7Networks_RH_SomMot_1 | 0.452 | 0.000 |
| 7Networks_RH_SomMot_2 | 0.286 | 0.000 |
| 7Networks_RH_SomMot_3 | 0.374 | 0.000 |
| 7Networks_RH_SomMot_4 | 0.316 | 0.000 |
| 7Networks_RH_SomMot_5 | 0.224 | 0.000 |
| 7Networks_RH_SomMot_6 | 0.319 | 0.000 |
| 7Networks_RH_SomMot_7 | 0.517 | 0.000 |
| 7Networks_RH_SomMot_8 | 0.287 | 0.000 |
| 7Networks_RH_SomMot_9 | 0.348 | 0.000 |
| 7Networks_RH_SomMot_10 | 0.202 | 0.000 |
| 7Networks_RH_SomMot_11 | 0.316 | 0.000 |
| 7Networks_RH_SomMot_12 | 0.440 | 0.000 |
| 7Networks_RH_SomMot_13 | 0.279 | 0.000 |
| 7Networks_RH_SomMot_14 | 0.480 | 0.000 |
| 7Networks_RH_SomMot_15 | 0.378 | 0.000 |

| | | |
|---|---|---|
| 7Networks_RH_SomMot_16 | 0.233 | 0.000 |
| 7Networks_RH_SomMot_17 | 0.481 | 0.000 |
| 7Networks_RH_SomMot_18 | 0.400 | 0.000 |
| 7Networks_RH_SomMot_19 | 0.369 | 0.000 |
| 7Networks_RH_DorsAttn_Post_1 | 0.273 | 0.000 |
| 7Networks_RH_DorsAttn_Post_2 | 0.192 | 0.000 |
| 7Networks_RH_DorsAttn_Post_3 | 0.274 | 0.000 |
| 7Networks_RH_DorsAttn_Post_4 | 0.189 | 0.001 |
| 7Networks_RH_DorsAttn_Post_5 | 0.226 | 0.000 |
| 7Networks_RH_DorsAttn_Post_6 | 0.395 | 0.000 |
| 7Networks_RH_DorsAttn_Post_7 | 0.208 | 0.000 |
| 7Networks_RH_DorsAttn_Post_8 | 0.397 | 0.000 |
| 7Networks_RH_DorsAttn_Post_9 | 0.359 | 0.000 |
| 7Networks_RH_DorsAttn_Post_10 | 0.378 | 0.000 |
| 7Networks_RH_DorsAttn_FEF_1 | 0.306 | 0.000 |
| 7Networks_RH_DorsAttn_FEF_2 | 0.269 | 0.000 |
| 7Networks_RH_DorsAttn_PrCv_1 | 0.283 | 0.000 |
| 7Networks_RH_SalVentAttn_TempOccPar_1 | 0.299 | 0.000 |
| 7Networks_RH_SalVentAttn_TempOccPar_2 | 0.240 | 0.000 |
| 7Networks_RH_SalVentAttn_TempOccPar_3 | 0.303 | 0.000 |
| 7Networks_RH_SalVentAttn_PrC_1 | 0.168 | 0.003 |
| 7Networks_RH_SalVentAttn_FrOperIns_1 | 0.381 | 0.000 |
| 7Networks_RH_SalVentAttn_FrOperIns_2 | 0.459 | 0.000 |
| 7Networks_RH_SalVentAttn_FrOperIns_3 | 0.308 | 0.000 |
| 7Networks_RH_SalVentAttn_FrOperIns_4 | 0.238 | 0.000 |
| 7Networks_RH_SalVentAttn_Med_1 | 0.211 | 0.000 |
| 7Networks_RH_SalVentAttn_Med_2 | 0.487 | 0.000 |
| 7Networks_RH_SalVentAttn_Med_3 | 0.381 | 0.000 |
| 7Networks_RH_Limbic_OFC_1 | 0.434 | 0.000 |
| 7Networks_RH_Limbic_OFC_2 | 0.335 | 0.000 |
| 7Networks_RH_Limbic_OFC_3 | 0.423 | 0.000 |
| 7Networks_RH_Limbic_TempPole_1 | 0.263 | 0.000 |
| 7Networks_RH_Limbic_TempPole_2 | 0.353 | 0.000 |
| 7Networks_RH_Limbic_TempPole_3 | 0.462 | 0.000 |
| 7Networks_RH_Cont_Par_1 | 0.196 | 0.000 |
| 7Networks_RH_Cont_Par_2 | 0.159 | 0.002 |
| 7Networks_RH_Cont_Par_3 | 0.198 | 0.000 |
| 7Networks_RH_Cont_Temp_1 | 0.246 | 0.000 |
| 7Networks_RH_Cont_PFCv_1 | 0.259 | 0.000 |
| 7Networks_RH_Cont_PFCl_1 | 0.276 | 0.000 |
| 7Networks_RH_Cont_PFCl_2 | 0.324 | 0.000 |
| 7Networks_RH_Cont_PFCl_3 | 0.385 | 0.000 |

| | | |
|---|---|---|
| 7Networks_RH_Cont_PFCl_4 | 0.427 | 0.000 |
| 7Networks_RH_Cont_PFCl_5 | 0.382 | 0.000 |
| 7Networks_RH_Cont_PFCl_6 | 0.336 | 0.000 |
| 7Networks_RH_Cont_PFCl_7 | 0.318 | 0.000 |
| 7Networks_RH_Cont_pCun_1 | 0.279 | 0.000 |
| 7Networks_RH_Cont_Cing_1 | 0.525 | 0.000 |
| 7Networks_RH_Cont_Cing_2 | 0.394 | 0.000 |
| 7Networks_RH_Cont_PFCmp_1 | 0.289 | 0.000 |
| 7Networks_RH_Cont_PFCmp_2 | 0.460 | 0.000 |
| 7Networks_RH_Default_Par_1 | 0.196 | 0.001 |
| 7Networks_RH_Default_Par_2 | 0.201 | 0.000 |
| 7Networks_RH_Default_Par_3 | 0.173 | 0.001 |
| 7Networks_RH_Default_Temp_1 | 0.331 | 0.000 |
| 7Networks_RH_Default_Temp_2 | 0.377 | 0.000 |
| 7Networks_RH_Default_Temp_3 | 0.463 | 0.000 |
| 7Networks_RH_Default_Temp_4 | 0.275 | 0.000 |
| 7Networks_RH_Default_Temp_5 | 0.214 | 0.000 |
| 7Networks_RH_Default_PFCv_1 | 0.403 | 0.000 |
| 7Networks_RH_Default_PFCdPFCm_1 | 0.349 | 0.000 |
| 7Networks_RH_Default_PFCdPFCm_2 | 0.331 | 0.000 |
| 7Networks_RH_Default_PFCdPFCm_3 | 0.361 | 0.000 |
| 7Networks_RH_Default_PFCdPFCm_4 | 0.481 | 0.000 |
| 7Networks_RH_Default_PFCdPFCm_5 | 0.498 | 0.000 |
| 7Networks_RH_Default_PFCdPFCm_6 | 0.402 | 0.000 |
| 7Networks_RH_Default_PFCdPFCm_7 | 0.280 | 0.000 |
| 7Networks_RH_Default_pCunPCC_1 | 0.432 | 0.000 |
| 7Networks_RH_Default_pCunPCC_2 | 0.435 | 0.000 |
| 7Networks_RH_Default_pCunPCC_3 | 0.254 | 0.000 |

***Supplementary Table 22.*** **Heritability of local surface area.**

| Parcel | $h^2$ | p |
|---|---|---|
| 7Networks_LH_Vis_1 | 0.610 | 0.000 |
| 7Networks_LH_Vis_2 | 0.554 | 0.000 |
| 7Networks_LH_Vis_3 | 0.399 | 0.000 |
| 7Networks_LH_Vis_4 | 0.406 | 0.000 |
| 7Networks_LH_Vis_5 | 0.331 | 0.000 |
| 7Networks_LH_Vis_6 | 0.619 | 0.000 |
| 7Networks_LH_Vis_7 | 0.645 | 0.000 |
| 7Networks_LH_Vis_8 | 0.398 | 0.000 |
| 7Networks_LH_Vis_9 | 0.472 | 0.000 |
| 7Networks_LH_Vis_10 | 0.795 | 0.000 |
| 7Networks_LH_Vis_11 | 0.447 | 0.000 |
| 7Networks_LH_Vis_12 | 0.591 | 0.000 |
| 7Networks_LH_Vis_13 | 0.644 | 0.000 |
| 7Networks_LH_Vis_14 | 0.462 | 0.000 |
| 7Networks_LH_SomMot_1 | 0.463 | 0.000 |
| 7Networks_LH_SomMot_2 | 0.509 | 0.000 |
| 7Networks_LH_SomMot_3 | 0.416 | 0.000 |
| 7Networks_LH_SomMot_4 | 0.345 | 0.000 |
| 7Networks_LH_SomMot_5 | 0.434 | 0.000 |
| 7Networks_LH_SomMot_6 | 0.498 | 0.000 |
| 7Networks_LH_SomMot_7 | 0.288 | 0.000 |
| 7Networks_LH_SomMot_8 | 0.477 | 0.000 |
| 7Networks_LH_SomMot_9 | 0.222 | 0.000 |
| 7Networks_LH_SomMot_10 | 0.363 | 0.000 |
| 7Networks_LH_SomMot_11 | 0.416 | 0.000 |
| 7Networks_LH_SomMot_12 | 0.479 | 0.000 |
| 7Networks_LH_SomMot_13 | 0.440 | 0.000 |
| 7Networks_LH_SomMot_14 | 0.291 | 0.000 |
| 7Networks_LH_SomMot_15 | 0.475 | 0.000 |
| 7Networks_LH_SomMot_16 | 0.380 | 0.000 |
| 7Networks_LH_DorsAttn_Post_1 | 0.407 | 0.000 |
| 7Networks_LH_DorsAttn_Post_2 | 0.405 | 0.000 |
| 7Networks_LH_DorsAttn_Post_3 | 0.483 | 0.000 |
| 7Networks_LH_DorsAttn_Post_4 | 0.390 | 0.000 |
| 7Networks_LH_DorsAttn_Post_5 | 0.246 | 0.000 |
| 7Networks_LH_DorsAttn_Post_6 | 0.213 | 0.000 |
| 7Networks_LH_DorsAttn_Post_7 | 0.354 | 0.000 |
| 7Networks_LH_DorsAttn_Post_8 | 0.322 | 0.000 |
| 7Networks_LH_DorsAttn_Post_9 | 0.417 | 0.000 |
| 7Networks_LH_DorsAttn_Post_10 | 0.364 | 0.000 |
| 7Networks_LH_DorsAttn_FEF_1 | 0.279 | 0.000 |
| 7Networks_LH_DorsAttn_FEF_2 | 0.244 | 0.000 |
| 7Networks_LH_DorsAttn_PrCv_1 | 0.330 | 0.000 |
| 7Networks_LH_SalVentAttn_ParOper_1 | 0.408 | 0.000 |
| 7Networks_LH_SalVentAttn_ParOper_2 | 0.470 | 0.000 |
| 7Networks_LH_SalVentAttn_ParOper_3 | 0.327 | 0.000 |
| 7Networks_LH_SalVentAttn_FrOperIns_1 | 0.619 | 0.000 |
| 7Networks_LH_SalVentAttn_FrOperIns_2 | 0.570 | 0.000 |
| 7Networks_LH_SalVentAttn_FrOperIns_3 | 0.528 | 0.000 |
| 7Networks_LH_SalVentAttn_FrOperIns_4 | 0.370 | 0.000 |

| | | | | | |
|---|---|---|---|---|---|
| 7Networks_LH_SalVentAttn_PFCl_1 | 0.314 | 0.000 | 7Networks_LH_Default_pCunPCC_2 | 0.494 | 0.000 |
| 7Networks_LH_SalVentAttn_Med_1 | 0.587 | 0.000 | 7Networks_LH_Default_pCunPCC_3 | 0.372 | 0.000 |
| 7Networks_LH_SalVentAttn_Med_2 | 0.435 | 0.000 | 7Networks_LH_Default_pCunPCC_4 | 0.435 | 0.000 |
| 7Networks_LH_SalVentAttn_Med_3 | 0.353 | 0.000 | 7Networks_LH_Default_PHC_1 | 0.602 | 0.000 |
| 7Networks_LH_Limbic_OFC_1 | 0.586 | 0.000 | 7Networks_RH_Vis_1 | 0.523 | 0.000 |
| 7Networks_LH_Limbic_OFC_2 | 0.590 | 0.000 | 7Networks_RH_Vis_2 | 0.492 | 0.000 |
| 7Networks_LH_Limbic_TempPole_1 | 0.548 | 0.000 | 7Networks_RH_Vis_3 | 0.459 | 0.000 |
| 7Networks_LH_Limbic_TempPole_2 | 0.459 | 0.000 | 7Networks_RH_Vis_4 | 0.519 | 0.000 |
| 7Networks_LH_Limbic_TempPole_3 | 0.423 | 0.000 | 7Networks_RH_Vis_5 | 0.238 | 0.000 |
| 7Networks_LH_Limbic_TempPole_4 | 0.603 | 0.000 | 7Networks_RH_Vis_6 | 0.646 | 0.000 |
| 7Networks_LH_Cont_Par_1 | 0.236 | 0.000 | 7Networks_RH_Vis_7 | 0.631 | 0.000 |
| 7Networks_LH_Cont_Par_2 | 0.246 | 0.000 | 7Networks_RH_Vis_8 | 0.369 | 0.000 |
| 7Networks_LH_Cont_Par_3 | 0.208 | 0.000 | 7Networks_RH_Vis_9 | 0.774 | 0.000 |
| 7Networks_LH_Cont_Temp_1 | 0.341 | 0.000 | 7Networks_RH_Vis_10 | 0.760 | 0.000 |
| 7Networks_LH_Cont_OFC_1 | 0.449 | 0.000 | 7Networks_RH_Vis_11 | 0.267 | 0.000 |
| 7Networks_LH_Cont_PFCl_1 | 0.446 | 0.000 | 7Networks_RH_Vis_12 | 0.578 | 0.000 |
| 7Networks_LH_Cont_PFCl_2 | 0.408 | 0.000 | 7Networks_RH_Vis_13 | 0.553 | 0.000 |
| 7Networks_LH_Cont_PFCl_3 | 0.386 | 0.000 | 7Networks_RH_Vis_14 | 0.401 | 0.000 |
| 7Networks_LH_Cont_PFCl_4 | 0.330 | 0.000 | 7Networks_RH_Vis_15 | 0.308 | 0.000 |
| 7Networks_LH_Cont_PFCl_5 | 0.226 | 0.000 | 7Networks_RH_SomMot_1 | 0.652 | 0.000 |
| 7Networks_LH_Cont_pCun_1 | 0.506 | 0.000 | 7Networks_RH_SomMot_2 | 0.568 | 0.000 |
| 7Networks_LH_Cont_Cing_1 | 0.516 | 0.000 | 7Networks_RH_SomMot_3 | 0.510 | 0.000 |
| 7Networks_LH_Cont_Cing_2 | 0.303 | 0.000 | 7Networks_RH_SomMot_4 | 0.513 | 0.000 |
| 7Networks_LH_Default_Temp_1 | 0.659 | 0.000 | 7Networks_RH_SomMot_5 | 0.317 | 0.000 |
| 7Networks_LH_Default_Temp_2 | 0.363 | 0.000 | 7Networks_RH_SomMot_6 | 0.508 | 0.000 |
| 7Networks_LH_Default_Temp_3 | 0.392 | 0.000 | 7Networks_RH_SomMot_7 | 0.428 | 0.000 |
| 7Networks_LH_Default_Temp_4 | 0.503 | 0.000 | 7Networks_RH_SomMot_8 | 0.314 | 0.000 |
| 7Networks_LH_Default_Temp_5 | 0.354 | 0.000 | 7Networks_RH_SomMot_9 | 0.213 | 0.000 |
| 7Networks_LH_Default_Par_1 | 0.336 | 0.000 | 7Networks_RH_SomMot_10 | 0.315 | 0.000 |
| 7Networks_LH_Default_Par_2 | 0.278 | 0.000 | 7Networks_RH_SomMot_11 | 0.520 | 0.000 |
| 7Networks_LH_Default_Par_3 | 0.160 | 0.002 | 7Networks_RH_SomMot_12 | 0.380 | 0.000 |
| 7Networks_LH_Default_Par_4 | 0.353 | 0.000 | 7Networks_RH_SomMot_13 | 0.368 | 0.000 |
| 7Networks_LH_Default_PFC_1 | 0.478 | 0.000 | 7Networks_RH_SomMot_14 | 0.460 | 0.000 |
| 7Networks_LH_Default_PFC_2 | 0.591 | 0.000 | 7Networks_RH_SomMot_15 | 0.204 | 0.000 |
| 7Networks_LH_Default_PFC_3 | 0.449 | 0.000 | 7Networks_RH_SomMot_16 | 0.302 | 0.000 |
| 7Networks_LH_Default_PFC_4 | 0.562 | 0.000 | 7Networks_RH_SomMot_17 | 0.160 | 0.003 |
| 7Networks_LH_Default_PFC_5 | 0.414 | 0.000 | 7Networks_RH_SomMot_18 | 0.475 | 0.000 |
| 7Networks_LH_Default_PFC_6 | 0.597 | 0.000 | 7Networks_RH_SomMot_19 | 0.444 | 0.000 |
| 7Networks_LH_Default_PFC_7 | 0.402 | 0.000 | 7Networks_RH_DorsAttn_Post_1 | 0.510 | 0.000 |
| 7Networks_LH_Default_PFC_8 | 0.446 | 0.000 | 7Networks_RH_DorsAttn_Post_2 | 0.339 | 0.000 |
| 7Networks_LH_Default_PFC_9 | 0.393 | 0.000 | 7Networks_RH_DorsAttn_Post_3 | 0.402 | 0.000 |
| 7Networks_LH_Default_PFC_10 | 0.447 | 0.000 | 7Networks_RH_DorsAttn_Post_4 | 0.232 | 0.000 |
| 7Networks_LH_Default_PFC_11 | 0.352 | 0.000 | 7Networks_RH_DorsAttn_Post_5 | 0.267 | 0.000 |
| 7Networks_LH_Default_PFC_12 | 0.333 | 0.000 | 7Networks_RH_DorsAttn_Post_6 | 0.247 | 0.000 |
| 7Networks_LH_Default_PFC_13 | 0.324 | 0.000 | 7Networks_RH_DorsAttn_Post_7 | 0.192 | 0.001 |
| 7Networks_LH_Default_pCunPCC_1 | 0.700 | 0.000 | 7Networks_RH_DorsAttn_Post_8 | 0.227 | 0.000 |

| | | |
|---|---|---|
| 7Networks_RH_DorsAttn_Post_9 | 0.451 | 0.000 |
| 7Networks_RH_DorsAttn_Post_10 | 0.393 | 0.000 |
| 7Networks_RH_DorsAttn_FEF_1 | 0.216 | 0.000 |
| 7Networks_RH_DorsAttn_FEF_2 | 0.586 | 0.000 |
| 7Networks_RH_DorsAttn_PrCv_1 | 0.337 | 0.000 |
| 7Networks_RH_SalVentAttn_TempOccPar_1 | 0.335 | 0.000 |
| 7Networks_RH_SalVentAttn_TempOccPar_2 | 0.222 | 0.000 |
| 7Networks_RH_SalVentAttn_TempOccPar_3 | 0.419 | 0.000 |
| 7Networks_RH_SalVentAttn_PrC_1 | 0.315 | 0.000 |
| 7Networks_RH_SalVentAttn_FrOperIns_1 | 0.584 | 0.000 |
| 7Networks_RH_SalVentAttn_FrOperIns_2 | 0.547 | 0.000 |
| 7Networks_RH_SalVentAttn_FrOperIns_3 | 0.435 | 0.000 |
| 7Networks_RH_SalVentAttn_FrOperIns_4 | 0.488 | 0.000 |
| 7Networks_RH_SalVentAttn_Med_1 | 0.577 | 0.000 |
| 7Networks_RH_SalVentAttn_Med_2 | 0.527 | 0.000 |
| 7Networks_RH_SalVentAttn_Med_3 | 0.488 | 0.000 |
| 7Networks_RH_Limbic_OFC_1 | 0.532 | 0.000 |
| 7Networks_RH_Limbic_OFC_2 | 0.516 | 0.000 |
| 7Networks_RH_Limbic_OFC_3 | 0.461 | 0.000 |
| 7Networks_RH_Limbic_TempPole_1 | 0.509 | 0.000 |
| 7Networks_RH_Limbic_TempPole_2 | 0.493 | 0.000 |
| 7Networks_RH_Limbic_TempPole_3 | 0.455 | 0.000 |
| 7Networks_RH_Cont_Par_1 | 0.283 | 0.000 |
| 7Networks_RH_Cont_Par_2 | 0.228 | 0.000 |
| 7Networks_RH_Cont_Par_3 | 0.225 | 0.000 |
| 7Networks_RH_Cont_Temp_1 | 0.370 | 0.000 |
| 7Networks_RH_Cont_PFCv_1 | 0.358 | 0.000 |
| 7Networks_RH_Cont_PFCl_1 | 0.527 | 0.000 |
| 7Networks_RH_Cont_PFCl_2 | 0.346 | 0.000 |
| 7Networks_RH_Cont_PFCl_3 | 0.396 | 0.000 |
| 7Networks_RH_Cont_PFCl_4 | 0.415 | 0.000 |
| 7Networks_RH_Cont_PFCl_5 | 0.327 | 0.000 |
| 7Networks_RH_Cont_PFCl_6 | 0.272 | 0.000 |
| 7Networks_RH_Cont_PFCl_7 | 0.418 | 0.000 |
| 7Networks_RH_Cont_pCun_1 | 0.416 | 0.000 |
| 7Networks_RH_Cont_Cing_1 | 0.469 | 0.000 |
| 7Networks_RH_Cont_Cing_2 | 0.379 | 0.000 |
| 7Networks_RH_Cont_PFCmp_1 | 0.569 | 0.000 |
| 7Networks_RH_Cont_PFCmp_2 | 0.425 | 0.000 |
| 7Networks_RH_Default_Par_1 | 0.314 | 0.000 |
| 7Networks_RH_Default_Par_2 | 0.206 | 0.000 |
| 7Networks_RH_Default_Par_3 | 0.198 | 0.000 |
| 7Networks_RH_Default_Temp_1 | 0.490 | 0.000 |
| 7Networks_RH_Default_Temp_2 | 0.416 | 0.000 |
| 7Networks_RH_Default_Temp_3 | 0.408 | 0.000 |
| 7Networks_RH_Default_Temp_4 | 0.374 | 0.000 |
| 7Networks_RH_Default_Temp_5 | 0.453 | 0.000 |
| 7Networks_RH_Default_PFCv_1 | 0.367 | 0.000 |
| 7Networks_RH_Default_PFCdPFCm_1 | 0.574 | 0.000 |
| 7Networks_RH_Default_PFCdPFCm_2 | 0.493 | 0.000 |
| 7Networks_RH_Default_PFCdPFCm_3 | 0.188 | 0.001 |
| 7Networks_RH_Default_PFCdPFCm_4 | 0.592 | 0.000 |
| 7Networks_RH_Default_PFCdPFCm_5 | 0.381 | 0.000 |
| 7Networks_RH_Default_PFCdPFCm_6 | 0.400 | 0.000 |
| 7Networks_RH_Default_PFCdPFCm_7 | 0.307 | 0.000 |
| 7Networks_RH_Default_pCunPCC_1 | 0.666 | 0.000 |
| 7Networks_RH_Default_pCunPCC_2 | 0.401 | 0.000 |
| 7Networks_RH_Default_pCunPCC_3 | 0.319 | 0.000 |

***Supplementary Table 23.*** **Heritability of subcortical volumes.**

| Volume | $h^2$ | p |
|---|---|---|
| accumb_l | 0.535 | 0.000 |
| accumb_r | 0.571 | 0.000 |
| amy_l | 0.621 | 0.000 |
| amy_r | 0.698 | 0.000 |
| caud_l | 0.837 | 0.000 |
| caud_r | 0.835 | 0.000 |
| hipp_l | 0.556 | 0.000 |
| hipp_r | 0.812 | 0.000 |

| | | |
|---|---|---|
| spall_l | 0.571 | 0.000 |
| pall_r | 0.666 | 0.000 |
| put_l | 0.709 | 0.000 |
| put_r | 0.848 | 0.000 |
| thal_l | 0.584 | 0.000 |
| thal_r | 0.667 | 0.000 |
| ventDC_l | 0.704 | 0.000 |
| ventDC_r | 0.718 | 0.000 |
